# Diffusion of the disordered E-cadherin tail on β-catenin

**DOI:** 10.1101/2021.02.03.429507

**Authors:** Felix Wiggers, Samuel Wohl, Artem Dubovetskyi, Gabriel Rosenblum, Wenwei Zheng, Hagen Hofmann

## Abstract

Specific protein interactions typically require well-shaped binding interfaces. Here, we report a cunning exception. The disordered tail of the cell-adhesion protein E-cadherin dynamically samples a large surface area of the proto-oncogene β-catenin. Single-molecule experiments and molecular simulations resolve these motions with high resolution in space and time. Contacts break and form within hundreds of microseconds without dissociation of the complex. A few persistent interactions provide specificity whereas unspecific contacts boost affinity. The energy landscape of this complex is rugged with many small barriers (3 – 4 *k*_B_*T*) and reconciles specificity, high affinity, and extreme disorder. Given the roles of β-catenin in cell-adhesion, signalling, and cancer, this Velcro-like design has the potential to tune the stability of the complex without requiring dissociation.

---

Specific molecular interactions orchestrate a multitude of simultaneous cellular processes. The discovery of intrinsically disordered proteins (IDPs)^1, 2^ has substantially aided our understanding of such interactions. More than two decades of research revealed a plethora of functions and mechanisms^2, 3, 4^ that complemented the prevalent structure-based view on protein interactions. Even the idea that IDPs always ought to fold upon binding has largely been dismantled by recent discoveries of high affine disordered complexes^5, 6^. Classical shape-complementary is indeed superfluous in the complex between prothymosin-α and histone H1 in which charge-complementary is the main driving-force for binding^5^. Yet, specificity remains limited in these complexes and it is unclear whether extreme levels of disorder, specificity, and affinity are compatible. In fact, many complexes between IDPs and folded proteins such as Sic1 and Cdc4^7^, the Na^+^/H^+^ exchanger tail and ERK2^8^, nucleoporin tails and nuclear transport receptors^9^ balance these factors. For instance, whereas nucleoporin tails are disordered in complex with Importin-β, affinities are weak and association/dissociation is fast^9^.

Here, we show that a cell adhesion complex involved in growth pathologies and cancer^10^ reconciles high affinity and specificity with flexibility. The intrinsically disordered cytoplasmic tail of E-cadherin retains disorder in complex with the multi-functional signalling factor β-catenin^11, 12, 13^. Large segments of E-cadherin diffuse dynamically on the β-catenin surface. By integrating single-molecule FRET (smFRET) experiments with molecular simulations, we obtained a residue-resolved understanding of these motions: A small number of persistent interactions provide specificity whereas many weak multivalent contacts boost affinity. With this design, regulatory enzymes can easily access their recognition motifs on E-cadherin and β-catenin whereas the elastic properties of the complex might dampen forces exerted by neighbouring epithelial cells.

E-cadherin is a trans-membrane protein that mediates cell-cell adhesions by linking actin filaments of adjacent epithelial cells (Fig. 1a). The cytoplasmic tail of E-cadherin binds β-catenin, which establishes a connection to the actin-associated protein α-catenin^14, 15, 16^. β-catenin on the other hand, is a multifunctional repeat protein^17, 18, 19, 20^ that mediates cadherin-based cell adhesions^21^ and governs cell fate decisions during embryogenesis^22^. It contains three domains: an N-terminal domain (130 aa), a central repeat domain (550 aa), and a C-terminal domain (100 aa). The 12 repeats of the central domain arrange in a super-helix^23^ and the X-ray structure shows that the E-cadherin tail (hereafter E-cad) wraps around the central domain of β-catenin (hereafter β-cat)^23^ (Fig. 1b). However, not only is half of the electron density of E-cad missing, the X-ray unit cell also comprises two structures with different resolved parts of E-cad (Fig. 1b). In fact, only 45% of all resolved E-cad residues are found in both structures (Fig. 1c). This ambiguity together with the large portion of missing residues might suggest^24^ that E-cad is highly dynamic in complex with β-cat.

**Fig. 1.**
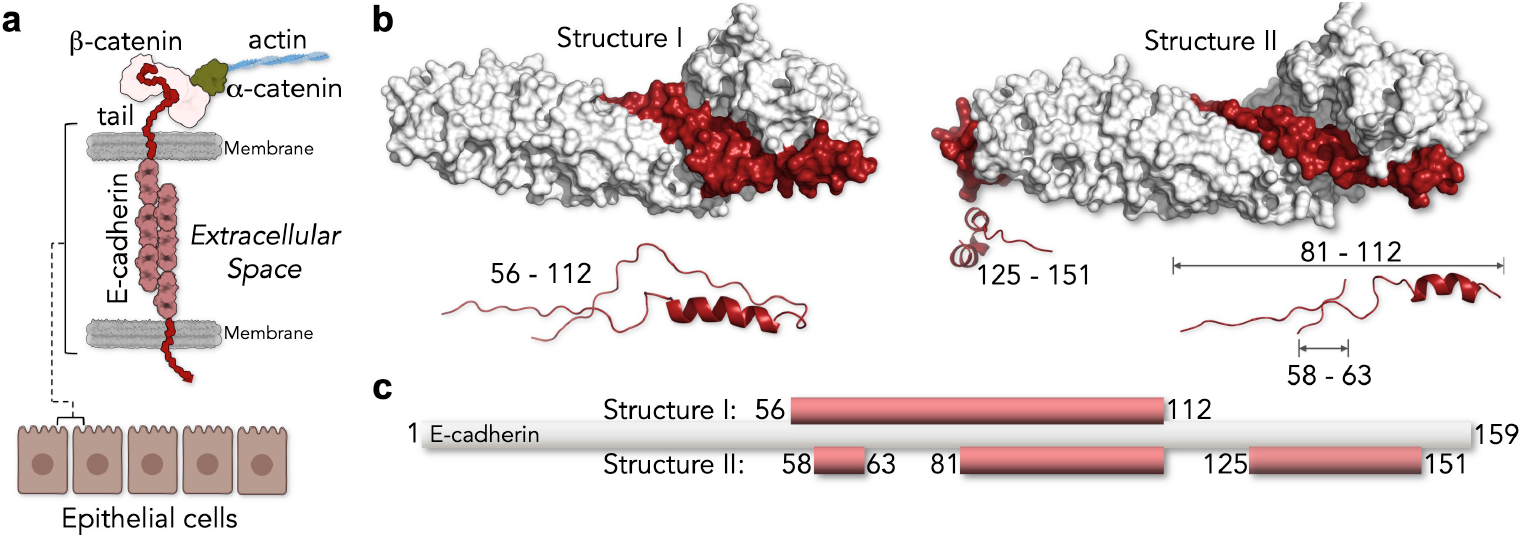
Complex between the cytoplasmic tail of E-cadherin (E-cad) and β-catenin (β-cat). (**a**) Schematics of cell-cell junctions mediated by E-cadherin and β-catenin. (**b**) The two X-ray structures of the complex between the tail of E-cadherin (red) and β-catenin (white) resolve different parts of E-cadherin (PDB: 1i7x). Bottom: Cartoon representation of the resolved E-cadherin parts. (**c**) Scheme showing the resolved parts of E-cadherin (red).

To understand how E-cad binds β-cat with high specificity and high affinity while retaining flexibility, we integrated smFRET experiments and molecular simulations. In our bottom-up strategy, we first probed intra-molecular interactions within E-cad using smFRET to parameterize a coarse-grained (CG) model. In a second step, we monitored E-cad on β-cat, integrated this information into the CG-model, and obtained a dynamic picture that reconciles specificity, affinity, and disorder.

## E-cad is expanded in solution

To probe the conformation of E-cad, we labelled different regions with AlexaFluor 488 as donor and AlexaFluor 594 as acceptor. We created three constructs in which the FRET labels map the N-terminal (A), central (B), and C-terminal (C) segments of E-cad (Fig. 2a, Supplementary Table 1). In addition, we also checked the longer segments AB, BC, and ABC to characterize the global behaviour of E-cad. The labelled E-cad constructs were monitored while freely diffusing through the confocal volume of our microscope. For all six constructs, we obtained homogeneous FRET distributions (Fig. 2a) with mean positions that scaled with the sequence separation between the dyes (Fig. 2b), as expected for an IDP. The persistence lengths of all segments, except of the A-segment, significantly exceeded the value of 0.4 nm found for other disordered or unfolded proteins^25^ (Fig. 2c). Hence, E-cad is expanded in solution, likely due to electrostatic repulsions given its high net charge (−22) ^26, 27, 28^. When we screened these repulsions with KCl, constructs ABC, AB, and BC collapsed substantially, a feature that is well described by mean-field polyampholyte theory^29^ (Fig. 2d). Yet, compaction was not uniform across the chain. Whereas the local segments B and C also compacted, the N-terminal A-segment expanded (Fig. 2d). Such chain expansion upon charge screening is known for polyampholytes with balanced numbers of positive and negative charges^28, 29, 30, 31, 32^. However, the A-segment is not charge-balanced. It even has a more negative net charge per residue (−0.186) than B (−0.093) and C (−0.166), thus excluding polyampholyte effects^29^. Instead, the A-segment exhibits significant charge segregation: a positively charged N-terminus is followed by a negative C-terminal part (Fig. 2a). Such charge patterning has previously been shown to compact disordered proteins^33^, which we confirmed using a sequence charge decoration metric (SCD_lowsalt_)^34, 35^ (Supplementary Fig. 1a). Mean-field polyampholyte theory is therefore inappropriate to describe E-cad, despite its success for other IDPs^28, 30^ (Fig. 2d). To convert our smFRET experiments into structural ensembles, we modelled E-cad as a coarse-grained (CG) heteropolymer of beads. Each bead corresponded to an amino acid with charge (+1, 0, −1) and hydrophobicity. The strength of hydrophobic interactions was adjustable via a global parameter *ε* (see Methods). Remarkably, a single value (*ε* = 0.16 kcal/mol) reproduced all experimental data, including those for the A-segment (Fig. 2d). A 2D-map of length scaling exponents (*ν*) for individual residue pairs^36^ now clearly uncovers the effect of charge segregation in the A-segment (Fig. 2e). The scaling exponents for residue pairs from the positively charged N-terminus and the negatively charged C-terminus of the A-segment are small (*ν* < 0.5), which suggested strong attractions between the termini. Swapping a pair of charged residues between the termini would therefore revert the salt-dependence of the A-segment (Supplementary Fig. 1b). Notably, high salt concentrations (>500 mM) collapsed all segments (Fig. 2d), a salting-out effect^30^ that is irrelevant at the physiological salt concentrations relevant for our study.

**Fig. 2.**
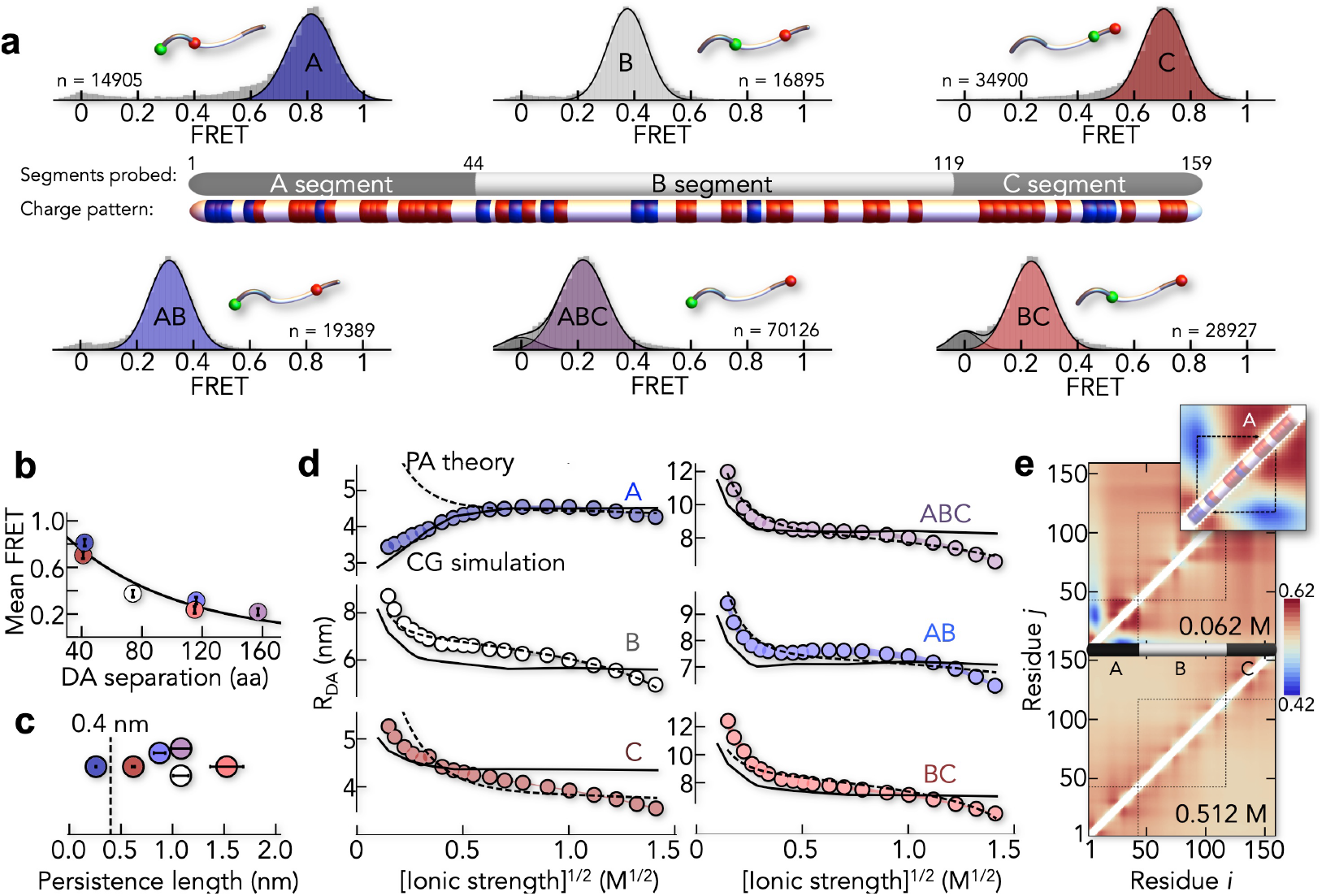
E-cad is expanded in solution. (**a**) SmFRET histograms of six segments of E-cad (A, B, C, AB, BC, ABC) with *n* indicating the total number of molecules. A scheme shows the division of E-cad into the segments probed by FRET and the distribution of positively (blue) and negatively charged (red) residues. Histogram insets indicate the labelling sites as green and red spheres for donor and acceptor, respectively. (**b**) The mean FRET efficiencies scale with the sequence separation of the dyes. The colour code is identical to (a). (**c**) Persistence lengths of the individual segments. Error bars are the uncertainty introduced by the model of the distance distribution used (see Methods). (**d**) Salt-induced change in the donor-acceptor distance *R_DA_* of the six E-cad segments (see Methods). A fit with the polyampholyte theory^28, 29^ (dashed line) and the best fit of the CG-model (solid line) are shown for comparison. (**e**) Length-scaling exponent map resulting from the best-fit CG-model at low (62 mM, upper panel) and high (512 mM, lower panel) ionic strength. Inset: Segment A.

## E-cad forms a pliable complex with β-cat

To monitor E-cad in the complex, we added unlabelled β-cat, which gave rise to a second population of E-cad molecules with altered FRET efficiencies (Fig. 3a). In most segments, the FRET efficiency was higher than in free E-cad, indicating a compaction upon binding to β-cat. An exception was again the A-segment. The disruption of interactions between its oppositely charged termini slightly expanded this segment upon binding to β-cat. We next performed smFRET experiments at different β-cat concentrations to determine the affinity of the complex (Fig. 3b), which we found to be in the range of 4 ± 2 nM (Fig. 3c), i.e., very high for all constructs. This agreed with previous measurements of unlabelled E-cad^37^, showing that labelling did not interfere with binding. Remarkably, whereas affinity was only moderately affected by salt (Fig. 3c), we found that increasing KCl concentrations caused a significant decrease in the FRET efficiencies of E-cad, indicative of an expansion (Fig. 3d). This solvent sensitivity suggested that E-cad in complex with *β*-cat retains significant flexibility.

**Fig. 3.**
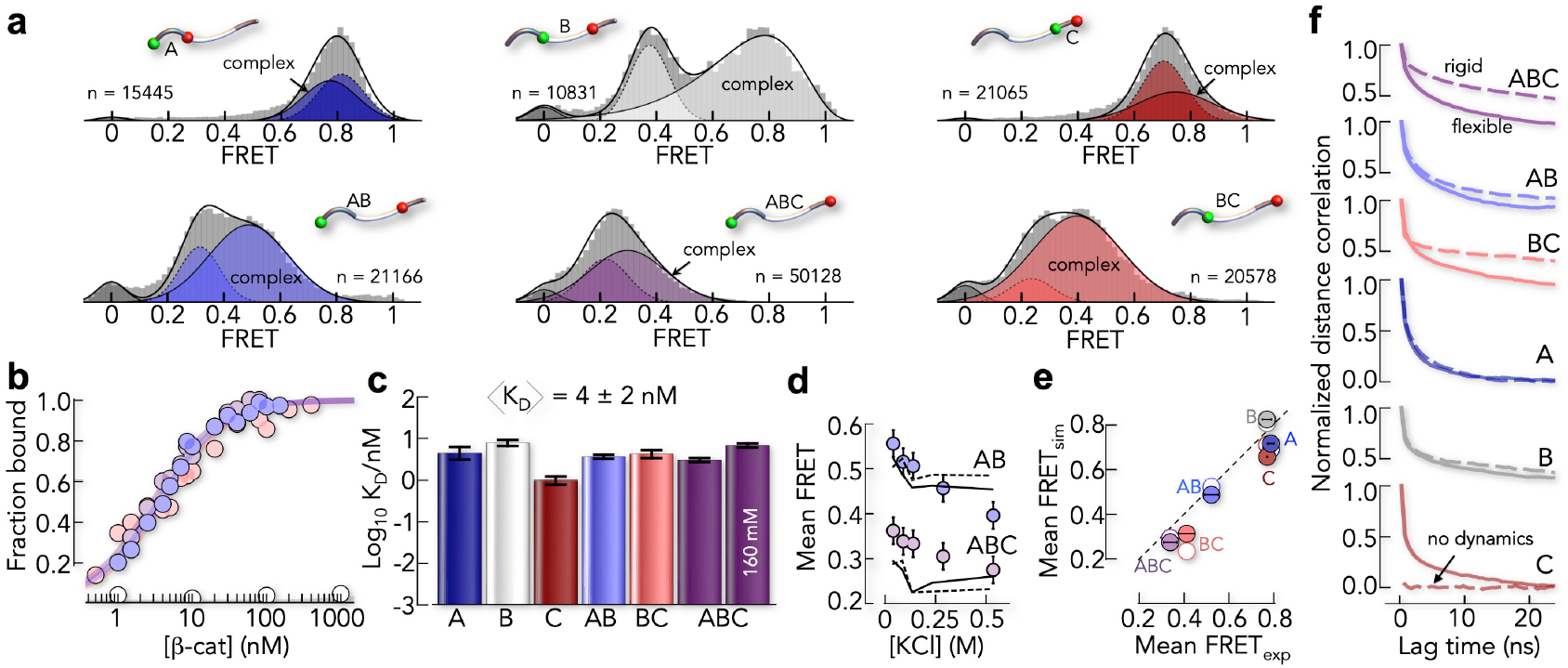
The conformation of E-cad in complex with β-cat is pliable. (**a**) smFRET histograms of the six E-cad constructs in the presence of β-cat (A: 4 nM, B: 85 nM, C: 15nM, AB: 7 nM, BC: 8.5 nM, ABC: 5 nM). Solid lines are fits of the histograms with a superposition of Gaussian and log-normal peaks and dashed lines represent free E-cad (see Methods). The number of molecules in each histogram is indicated by *n*. (**b**) Binding isotherms of selected E-cad segments (filled circles) and of the core-binding peptide (white circles). (**c**) Dissociation constants (K_D_) for the six constructs at 82 mM ionic strength and a comparison of the affinity of the ABC segment at higher ionic strength. (**d**) Change in FRET efficiency of the E-cad constructs ABC and AB in complex with β-cat with increasing ion concentrations. Circles indicate the experimental data and lines are the result from the rigid CG-model when keeping the X-ray resolved E-cad parts static (dashed) and the best fit of the flexible CG-model. (**e**) Comparison of the experimental FRET efficiencies with the rigid CG-model (open circles) and the flexible CG-model (filled circles). (**f**) Donor-acceptor distance correlation functions from the rigid CG-model (dashed lines) and the flexible CG-model (solid lines).

To understand this flexibility, we extended our CG-model by intermolecular contacts between E-cad and β-cat (see Methods). We first tested a CG-model in which we rigidified the parts of E-cad that are resolved in the X-ray structures^23^. Hence, structured parts of E-cad were essentially kept static while the unresolved parts were flexible but allowed to contact β-cat. Indeed, this ‘rigid’ model described the observed FRET efficiencies (Fig. 3e). The model even partially captured the experimentally observed expansion of E-cad at higher salt concentrations (Fig. 3d), suggesting that flexible parts are responsible for the effect. How flexible are the individual segments?

Our model predicted extensive motions in all segments except the C-segment, which interacted rigidly with β-cat (Fig. 3f). Unfortunately, simplifications inherent to coarse-grained models prevented accurate timescale estimates. We therefore measured the dynamics of E-cad in complex with β-cat experimentally.

## E-cad is highly dynamic in complex with β-cat

To probe the dynamics of the E-cad ensemble, we performed nanosecond FCS (nsFCS)^38, 39^ experiments of labelled E-cad in the absence and presence of β-cat. Disordered proteins such as E-cad exhibit large-scale distance fluctuations that cause anti-correlated donor-acceptor intensity fluctuations^39^. Indeed, the cross-correlation functions of free E-cad reveal anti-correlated decays in the 100 ns timescale for almost all segments (Fig. 4a). Only the C-segment lacks this signal, likely due to inter-dye contacts caused by a close proximity of the dyes in this short segment^40^ (Supplementary Fig. 2a).

**Fig. 4.**
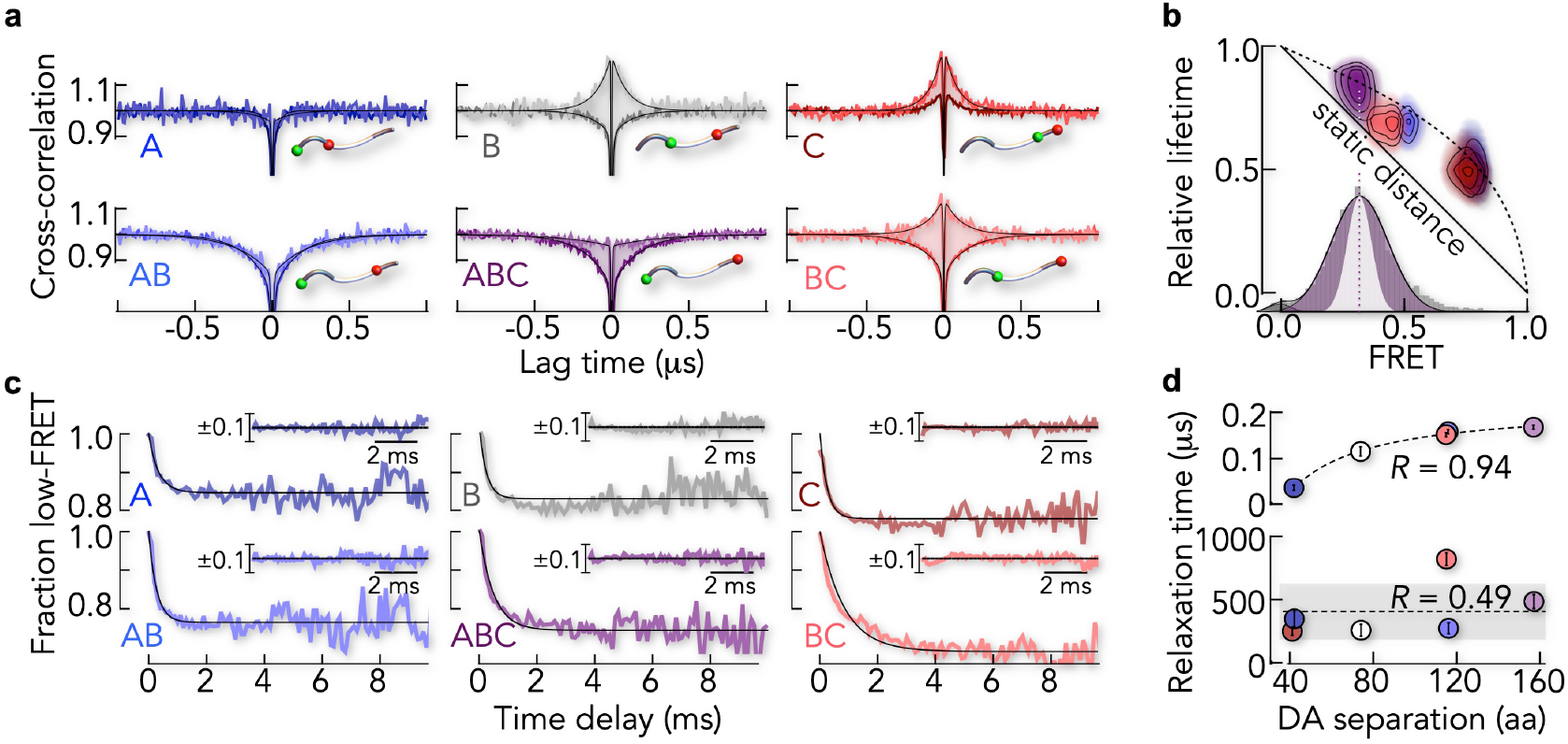
Timescales of motion of E-cad free and bound to β-cat. (**a**) Donor-acceptor cross-correlation functions of all six FRET constructs of E-cad in the absence (darker colour) and presence (lighter colour) of saturating amounts of β-cat (100 nM). Solid lines are fits to a product of exponential terms (see Methods). (**b**) 2D correlation map between the donor fluorescence lifetime and FRET efficiency of all six E-cad constructs in complex with β-cat. The solid line shows the expected dependence for a single donor-acceptor distance. The dashed line indicates the dependence for a Gaussian chain distribution as an upper limit of distance heterogeneity. Inset: Measured FRET histogram of the ABC segment of E-cad in complex with β-cat in comparison to the expected shot-noise limit (white area). (**c**) RASP time decays of the fraction of molecules with FRET values lower than the mean value of the FRET histogram for all six E-cad contructs in complex with β-cat. Solid lines are exponential fits of the data. Inset: residuals of the exponential fit. (**d**) Comparison of the relaxation times of free E-cad (upper panel) and β-cat bound E-cad (lower panel) as a function of the donor-acceptor sequence separation. The Pearson correlation coefficient is indicated.

When we formed the complex by adding β-cat, the correlation functions changed. Whereas signals in A and AB were nearly unaltered, the cross-correlation amplitude of the ABC segment decreased, indicating reduced fluctuations at the sub-microsecond timescale (Fig. 4a). Segments B, C, and BC even showed positive cross-correlation amplitudes, thus providing no information about potential distance dynamics. Dye-sticking to protein surfaces had previously been reported as cause for such positive cross-correlation amplitudes^41^, but we excluded this possibility with fluorescence anisotropy experiments (Supplementary Fig. 3). Instead, we found increased static quenching, likely due to dye contacts with aromatic residues of β-cat (Supplementary Fig. 2b).

To bypass this effect, we determined donor fluorescence lifetimes for all segments, a quantity that is unaffected by static quenching. For a static well-defined distance, these lifetimes are expected to correlate linearly with the FRET efficiency^39^ (Fig. 4b). Yet, in contrast to this prediction, we found substantial deviations from the expected values, suggesting an ensemble of structures. Importantly, we found this deviation also for the C-segment (Fig. 4b). This is surprising given that the C-segment is resolved in the X-ray structure (Fig. 1b), did not show flexibility in our nsFCS experiments (Fig. 4a) and therefore was not expected to be dynamic. However, nsFCS experiments only probe dynamics on sub-microsecond timescales but cannot capture slower motions. In fact, we observed substantial broadening of the FRET peaks of all segments in complex with β-cat (Fig. 4b inset), which is symptomatic for dynamics in the millisecond regime.

We can readily identify such slow dynamics using RASP (recurrence analysis of single particles)^42, 43, 44^. The method exploits the fact that a diffusing molecule can enter and exit the confocal volume multiple times. Once a molecule leaves the observation volume, the chance of it returning within a short time interval is greater than the chance of detecting a new molecule. This effect allowed us to monitor conformational switching between the low-FRET (left) and high-FRET (right) side of the FRET distributions (see Methods, Supplementary Fig. 4). The resulting exchange kinetics indeed showed pronounced decays on timescales of micro- to millisecond for all six constructs (Fig. 4c), including the C-segment. The relaxation times ranged from 250 ± 30 μs for the C-segment to 820 ± 50 μs for the BC-segment, i.e., three orders of magnitude slower than for free E-cad. Moreover, the relaxation times did not scale with the sequence separation of the dyes as expected for polymers^45, 46, 47^ such as free E-cad (Fig. 4d). This showed that motions were determined by contacts between E-cad and β-cat rather than within E-cad. Most importantly, these experiments clearly showed that even the C-segment, which was expected to be rigidly bound to β-cat, is highly dynamic. How can we reconcile this finding with the X-ray structure? In fact, the C-segment is only resolved in one of two structures in the unit cell (Fig. 1b,c). Such ambiguity has also been observed in other complexes^24^ and suggests that the C-segment can indeed dissociate from the β-cat surface, which explains the dynamics found experimentally. Hence, our results show that the E-cad/β-cat complex is more dynamic than indicated by the X-ray structure and our rigid CG-model.

## A structural model of E-cad in complex with β-cat

To obtain a more realistic picture of the E-cad ensemble in complex with β-cat, we revised our CG-model. Instead of fully rigidifying the resolved parts of E-cad, we introduced a parameter (ζ) that tunes the strength of all intermolecular X-ray contacts in a global manner. We found that ζ = 0.6 kcal/mol described the experimentally determined FRET-values and the salt-sensitivity of the complex equally good as the rigid model (Fig. 3d-e). Most importantly, despite the fact that all X-ray contacts share the same interaction parameter ζ, the model indeed predicted a dynamic C-segment (Fig. 3f), in accord with our experimental findings.

To understand the deviations of this model from the known X-ray structure, we compared the distribution of contact types. To this end, we classified amino acids as polar (P), charged (C), hydrophobic (H), and other amino acids (O). Interestingly, the most abundant intermolecular contacts in the CG-model were of the uncommon H-C type instead of the commonly discussed C-C type (Fig. 5a), which explains the weak salt-dependence of the binding affinity (Fig. 3c). However, this distribution of contact types agrees well with the X-ray structure (Fig. 5a) and an intermolecular contact map shows that interactions found in the X-ray structure were also preserved in the CG-model (Fig. 5b). Yet, the smFRET-based CG-model differed from the known structure in one important aspect. Contact probabilities (Fig. 5b) and lifetimes (Supplementary Fig. 5) were broadly distributed in the CG-model, reflecting many weak and unspecific interactions. The highest contact probabilities, i.e., the most persistent interactions, were found in a short 20 amino acid stretch of the B-segment (Fig. 5b), a region that is well resolved in both X-ray structures (Fig. 1b,c). It has previously been identified as core-binding region of E-cad^48^ and likely provides specificity. To disentangle contributions of specific and unspecific interactions to the overall stability of the E-cad/β-cat complex, we determined its affinity to β-cat (Fig. 3b). We found a low affinity of the core-binding region with an upper limit of *K_D_* > 1 μM compared to *K_D_* ~ 4 nM for full-length E-cad. Hence, flexible segments of E-cad boost affinity by more than 5.5 *k*_B_*T*. How are these segments distributed on the surface of β-cat?

**Fig. 5.**
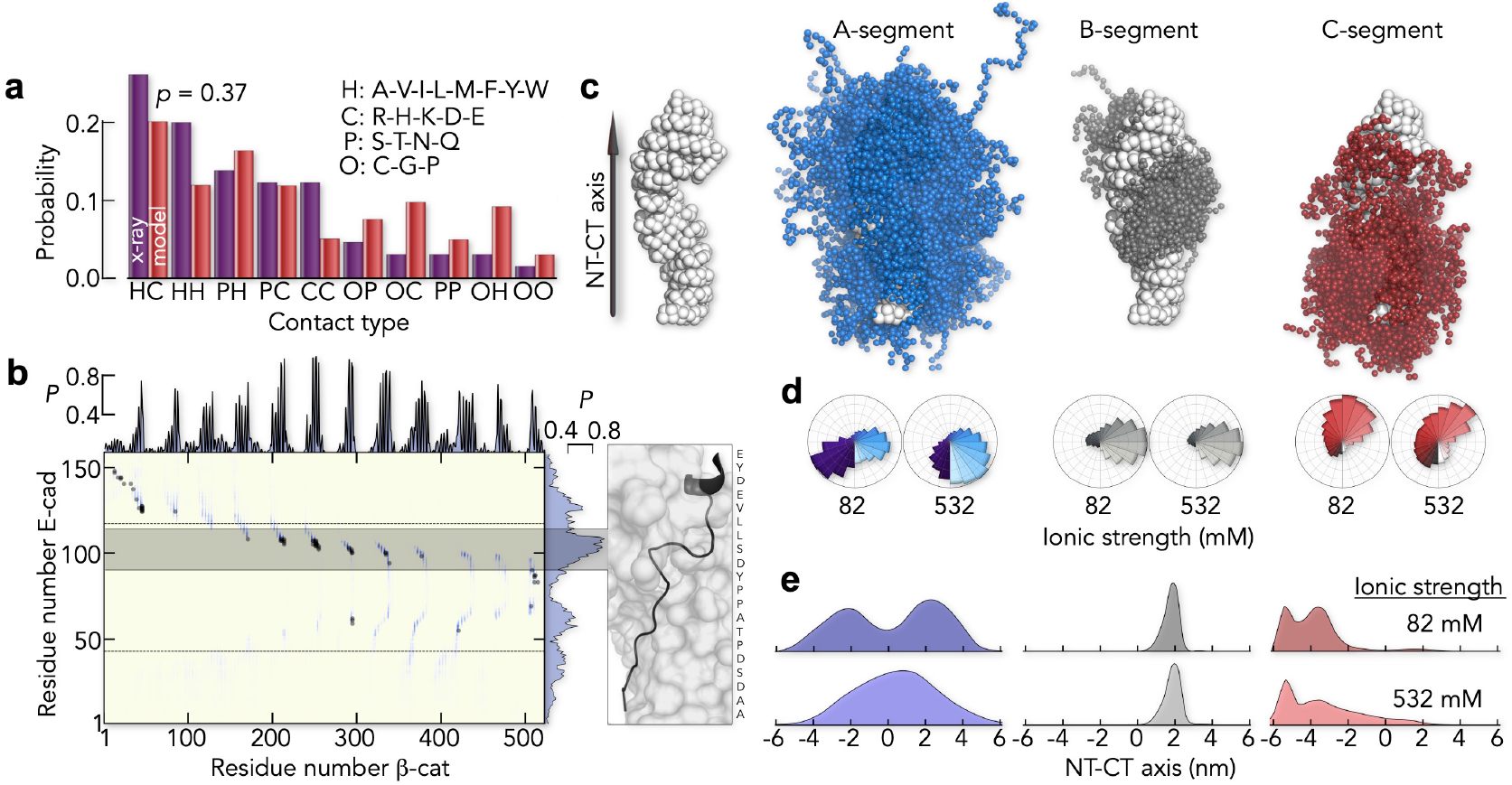
The flexible CG-model predicts a heterogeneous structural ensemble of E-cad in complex with β-cat. (**a**) Distribution of contact types in the complex. The classification of contacts is indicated. A Komolgorov-Smirnov test gives a probability of *p* = 0.37 that both distributions originate from the same sample. (**b**) Intermolecular contact probabilities (*P*) based on the flexible CG-model (blue) overlaid with the contact map derived from the X-ray map (gray dots). The highest contact probabilities are found in a 20 amino acid long fragment in the B-segment (gray bar) that is well resolved in both X-ray structures (right panel). (**c**) Structure of β-cat indicating the directionality of the axis between (left) and ensemble of the centre of mass positions (right) of A (blue), B (gray), and C (red) segment. (**d**) Centre of mass angular distribution along the axis between the N-terminus (NT) and C-terminus (CT) of β-cat (schematically indicated) at low (82 mM) and high (532 mM) ionic strength. Colour code is identical to (c). (**e**) Centre of mass positional distribution of E-cad projected onto the NT-CT axis of β-cat for the segments A, B, and C at two ionic strengths (indicated). The colour code is identical to (d) and (c).

The three-dimensional distribution of E-cad on β-cat differed strongly between segments (Fig. 5c). For instance, not only did the A-segment contact the N-terminus (NT) and the C-terminus (CT) of β-cat, but also the C-segment explored a large surface area of β-cat. The ensemble of the B-segment on the other hand was much more confined, as expected from the higher contact probabilities and the fact that 42% of it is resolved in both X-ray structures (Fig. 1c). We visualized the ensembles of the three segments by the distribution of their centre-of-mass along two coordinates: an angular coordinate describing the distribution around the long axis of β-cat (Fig. 5d), and an axial coordinate for the distribution along β-cat (Fig. 5e). These distributions showed that the observed salt-induced changes of E-cad in complex with β-cat (Fig. 3d) were predominantly due to changes in the A-segment (Fig. 5e,d) whereas changes in the B- and C-segment were much less pronounced.

In summary, the refined flexible CG-model indeed recapitulated the experimentally observed large-scale motions of E-cad on the β-cat surface. A total of 91% of the β-cat surface was explored by E-cad in a rather diffusive, i.e., continuous, manner without encountering major barriers. However, given the simplification inherent to CG-models, it remains to be determined whether a continuous diffusion of E-cad on the β-cat surface is realistic. An alternative model requires the concerted association and dissociation of E-cad segments, thus leading to dominant barriers that must be overcome to reconfigure on the β-cat surface. In fact, such a scenario would explain why E-cad motions in the complex (Fig. 4c) are orders of magnitude slower than those found for free E-cad (Fig. 4a). We therefore aimed at identifying the magnitude of these barriers.

## Intra-chain diffusion of E-cad on β-cat

The experimental RASP-kinetics of the six constructs agreed with exponential decays (Fig. 4c), a hallmark of two-state kinetics, and indicative of a dominant barrier. We tested this hypothesis by monitoring the motions of E-cad ABC at different temperatures (Supplementary Fig. 6). A barrier should cause Arrhenius behaviour, i.e., a significant slow-down of the dynamics with decreasing temperature. Yet, we only found a twofold slow-down of the kinetics in the range from 37°C to 5°C (Fig. 6a). A fit with the Arrhenius equation *τ* = *A* exp (*βE*_*a*_) with *β*^−1^ = *k_B_T* yielded an activation energy of *E_a_* = 28 ± 16 kJ/mol (Fig. 6b). However, this analysis neglected that the pre-exponential factor *A* depends on the viscosity of the medium^49^, which itself scales with temperature.

**Fig. 6.**
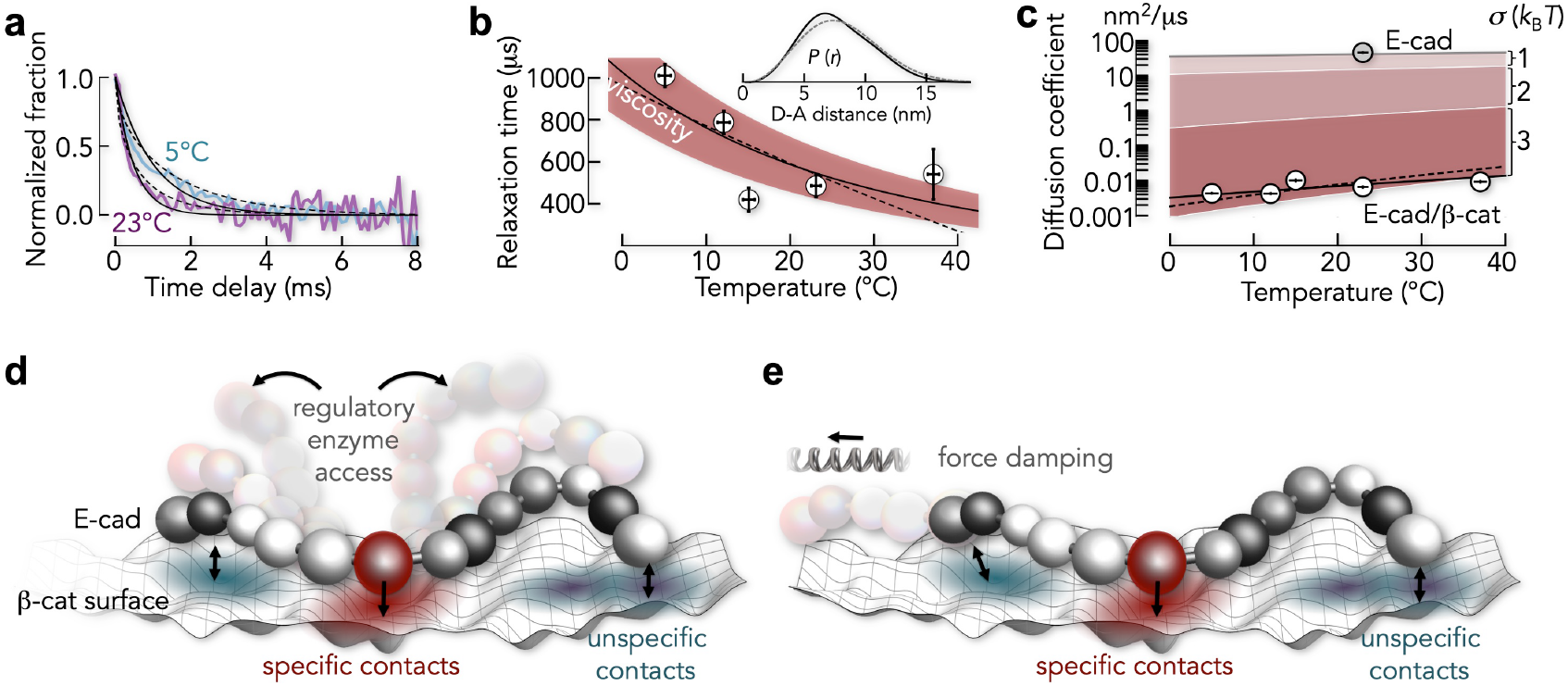
E-cad dynamics in complex with β-cat are diffusive. (**a**) Normalized time decays of the E-cad ABC construct at two temperatures (indicated). Solid and dashed lines are fits with an exponential function and with the Smoluchowski equation, respectively. (**b**) Relaxation times of the E-cad ABC construct as function of temperature. The solid line is the dependence without activation energy (see Methods). The red area indicates the 90% confidence interval of this fit. The dashed line is an Arrhenius fit. Inset: Donor-acceptor distance distributions for E-cad bound (black) and free (gray) from smFRET-based CG-model. (**c**) Comparison of the diffusion coefficient of E-cad free (gray circle) and bound to β-cat (white circles). Shaded areas represent the effect of different roughness values (σ in *k*_B_*T* at 23°C). Black solid and dashed lines are fits with the Zwanzig expressions^50^ for periodic (σ = 4.3 ± 0.1) and Gaussian (σ = 2.9 ± 0.1) modulated roughness, respectively. (**d**) Scheme of E-cad on the surface of β-cat. Parts of E-cad can partially detach, thus giving access to regulatory enzymes. (**e**) Same as d but illustrating the spring-like properties of the E-cad/β-cat complex.

Accounting for this effect, we found that the slow-down was also reproduced without activation energy (Fig. 6b). We therefore checked whether the kinetics were indeed consistent with single-well, i.e., barrier-less diffusion as suggested by our CG-model. To this end, we used the donor-acceptor distance distribution ***P*(*r*)** for the ABC construct from our flexible CG-model (Fig. 6b inset) to fit the experimental kinetics by solving the Smoluchowsky equation for diffusion in a single-well potential given by ***V*(*r*)** = − ln ***P*(*r*)**. Indeed, this barrier-less model also resulted in excellent fits. Intra-chain diffusion coefficients ranged from (4.4 ± 0.1)×10^−3^ nm^2^/μs to (9.5 ± 0.1)×10^−3^ nm^2^/μs in the temperature range from 5°C to 37°C (Fig. 6c, see Methods). Notably, intra-chain diffusion was four orders of magnitude slower than that of free E-cad ABC (42 ± 1 nm^2^/μs, see Methods) despite the absence of a dominant barrier (Fig. 6c).

Ruggedness in energy landscapes, i.e., a plethora of small barriers, is known to slow down motions as effectively as a dominant barrier^50, 51, 52, 53^. Zwanzig^50^ derived expressions for the apparent diffusion coefficient (*D*\*) (i) for randomly distributed barriers with average height σ, leading to 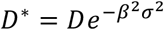 and (ii) for a periodic modulation, leading to ***D***\* = ***De***^−2*βσ*^. Here, *D* and *D*\* are diffusion coefficients in the absence and presence of ruggedness, respectively. Experimentally, it has been challenging to obtain absolute values for ***σ*** because *D* and *D*\* are typically not simultaneously known. Our experiments allowed us to overcome this problem. Since the donor-acceptor distance distribution of E-cad ABC changed only slightly upon binding to β-cat (Fig. 6b inset), the free energy potentials obtained from the smFRET-based CG-model were very similar between free and bound E-cad. Hence, *D* and *D*\* were simply the diffusion coefficients of isolated E-cad and bound to β-cat, respectively. Using the above relationships, we found the amplitude to be 2.9 ± 0.1 and 4.3 ± 0.1 (in *k*_B_*T* for *T* = 23°C) for randomly and periodically fluctuating ruggedness, respectively. These numbers suggest that the individual E-cad segments diffuse on a frustrated energy landscape with many local barriers of height 3 − 4 *k*_B_*T*.

## Conclusion

Whereas disordered complexes, sometimes being referred to as ‘fuzzy’, have been identified in the past^54^, their dynamics were often only accessible in simulations^55^. Our measurements of such dynamics in the E-cad/β-cat complex uncovered a rugged energy landscape of E-cad with many shallow minima. The minima depth, i.e., the total strength of all contacts is in the order of a hydrogen bond^56^, which is insufficient to rigidify E-cad. Nevertheless, the flexible parts of E-cad serve an important function: they boost affinity. This is necessary because specific contacts of the core-binding region fail to generate sufficient binding strength as indicated by the low affinity of the core-binding region of E-cad (Fig. 3b). Yet, the correct register of this element in the binding groove of β-cat provides specificity, which sets this complex apart from fully disordered complexes^5, 6^.

What might be the functional role of this unusual design? In-cell concentrations of β-cat are very low (< 1 nM)^57, 58^, even in adherent cancer cell lines where β-cat is overexpressed. Given the nano-molar affinity of the complex, adherens junctions would be unstable. Post-translational modifications of E-cad are known to increase the stability of the E-cad/β-cat complex^59^. However, allowing kinases to access their substrate by dissociating the entire complex requires hours for re-associating it with the known rates (10^5^ – 10^6^ M^−1^s^−1^)^37^. The continuous attachment and detachment of local E-cad segments bypasses this problem by providing easy access for regulatory enzymes without dissociating and re-associating the complex (Fig. 6d). In addition, the complex can also serve mechanical functions. E-cad and β-cat link the cytoskeletons of neighbouring epithelial cells (Fig. 1a), suggesting that rupture forces can propagate from cell to cell. Like all polymers, IDPs are soft entropic springs. Many weak contacts to the β-cat surface can increase the spring constant without causing a rigid complex (see Methods). The E-cad/β-cat complex has therefore the potential to act as a tuneable damper for mechanical forces between cells.

The Velcro-like design of many weak contacts on top of a few persistent interactions reconciles three seemingly contradictory factors: specificity, high affinity, and flexibility. Given the simplicity and advantages of this design together with the fact that partially disordered complexes are highly abundant^54^, the mode of interaction described here may be common in biology.

## Methods

### Constructs

The plasmid containing the cytoplasmic domain of mouse E-cad (residues 734-884, UniProt ID P09803) was a kind gift of William I. Weis from Stanford University School of Medicine. E-cad was transferred from the pGEX vector to a K151 vector system using transfer-PCR. The procedure was carried out as described previously^60^. Additionally, cysteine linkers were incorporated in the E-cad-K151 vector using the restriction free cloning approach^61^ at different positions to generate the six labelling variants (Table S1). Full-length murine-beta-catenin (UniProt ID Q02284, purchased from GenScript) was transferred to a K151 vector system using transfer-PCR. Both proteins start with an N-terminal His_12_-tag followed by the Sumo protein.

### Protein expression

Proteins were expressed in *E. coli* BL21(DE3) cells. A 50 ml overnight culture grown at 37 °C (100 μg ml^−1^ Kanamycin) was used to inoculate 4 l of super broth (100 μg ml^−1^ Kanamycin). The cultures were grown at 37 °C to an optical density of 0.6 - 0.8 cm^−1^ (at 600 nm). Then 0.5 mM isopropyl-β-D-thiogalactoside was added to induce protein expression for 3 h. After induction, the cultures were cooled to 4 °C and centrifuged at 3000 g for 30 min at 4 °C. The cell pellet was washed with 80 ml of ice-cold buffer (10 mM Tris-HCl pH 8.0, 300 mM NaCl) and centrifuged at 3000 g for 30 min at 4 °C. The supernatants were discarded and the pellets were stored at −80 °C.

### Protein purification

The cell pellet from 2 l culture was re-suspended in 50 mM Tris-HCl pH 8.0, 100 mM NaCl, 10 % glycerol, 1 mM DTT, 0.5 mM phenyl-methanesulfonyl fluoride (PMSF), 5 μM leupeptin, 2.5 μg ml-1 pepstatin (25 ml per 5 g wet weight) followed by sonication on ice using an ultrasonicator (Vibra-Cell, Sonics) at 70 % amplitude for 5 times 30 s with a 2 s on / 8 s off pulse. The soluble fraction was collected by centrifugation at 40,000 g for 30 min at 4 °C. Following sonication, DNA impurities were digested (DNase: Fisher Optizyme, 2 U/ml) and the supernatant was filtered (0.45 μm, Sartorius) and loaded at 4 °C with a flow rate of 0.5 ml min^−1^ onto a 5 ml HisTrap HP column (GE Healthcare) equilibrated with 50 mM Tris-HCl pH 8.0, 100 mM NaCl, 10 % glycerol. The column was subsequently washed with 20 ml of the equilibration buffer followed by 20 ml of 50 mM Tris-HCl pH 8.0, 500 mM NaCl and with additional 20 ml of 50 mM Tris-HCl pH 8.0, 470 mM NaCl, 75 mM imidazole at a flow rate of 1 ml min^−1^. The protein was eluted with 20 ml of 50 mM Tris-HCl pH 8.0, 300 mM NaCl, 500 mM imidazole at a flow rate of 1 ml min^−1^. Protein fractions were pooled.

### Removal of the His_6_-Sumo tag

The affinity tags of both proteins were removed using Sumo protease, bdSENP1^62^. The plasmid was a kind gift of Professor Dirk Görlich, Max-Planck-Institute for Biophysical Chemistry (Göttingen, Germany). A solution containing 300 nM Sumo protease was dialyzed twice against 50 mM Tris-HCl pH 8.0, 500 mM NaCl for 2h at 4°C. Afterwards, the sample was loaded at 2 ml min^−1^ onto a 5 ml HisTrap HP column (GE Healthcare) equilibrated with 50 mM Tris-HCl pH 8.0, 470 mM NaCl, 75 mM imidazole at 4°C. The column was washed with 20 ml of the same buffer. The flow-through containing the cleaved protein was collected. The protease and non-cleaved proteins were eluted with 20 ml of buffer B at a flow rate of 2 ml min^−1^. Protein fractions were pooled and submitted to size exclusion chromatography (SEC) using a Superdex 200 pg HiLoad 26/600 column (GE Healthcare Life Sciences) equilibrated with 20 mM Tris/HCl, 300 mM NaCl, 5 mM DTT, pH 8.0. Protein fractions were collected and pooled. Final refinement of the β-cat was conducted with a buffer exchange using a 5 ml GE, HiTrap desalting column into 25 mM Tris/HCl, 50 mM KCl, 200 mM L-Arg, pH 8, followed by concentrating the sample (VivaSpin 6 MWCO 3 kDa cut-off, GE Healthcare) to 8 μM. The E-cad constructs were concentrated to ~ 14 μM. Protein concentrations were determined based on the absorbance at 280 nm using the extinction coefficient at 280 nm of 18910 M^−1^cm^−1^ for the E-cad variants and 63830 M^−1^cm^−1^ for β-cat. All E-cad constructs were additionally purified using reverse-phase HPLC purification with the ZORBAX Eclipse Plus C18 (3.5 μm) column (Agilent) (see following section). The correct protein mass was confirmed via ESI mass spectrometry analysis. The proteins were drop-frozen in liquid nitrogen and stored at −80 °C.

### Fluorophore labeling with AlexaFluor 488 (donor) and AlexaFluor 594 (acceptor)

The six E-cad variants were reduced before labelling by the addition of 100 mM DTT. After 15 min at room temperature, the proteins were purified via reversed phase HPLC using a ZORBAX Eclipse Plus C18 (3.5 μm) column (Agilent) equilibrated with 0.1% trifluoroacetic acid (TFA). The E-cad constructs were eluted with gradients from 15% - 60% acetonitrile (ACN) in 13 ml. Purified proteins were lyophilized and stored at −80 °C. The lyophilized proteins were re-solubilized in 50 mM sodium phosphate pH 7.3, 6 M GdmCl at a protein concentration of 150 μM. The precise concentrations were determined using the absorbance at 280 nm. For labelling, the proteins were incubated with 0.6 equivalents of Alexa 488 C5 maleimide (Invitrogen) for 60 min (25 °C, 300 rpm). Afterwards, a tenfold excess of β-mercaptoethanol (compared to the dye) was added to quench unreacted dye. Reversed-phase HPLC was then used to remove unreacted dye, unlabelled and double-donor labeled protein. To this end, a ZORBAX Eclipse Plus C18 (3.5 μm) column (Agilent) equilibrated with water containing 0.1 % TFA was used. The gradient was 38% - 46% ACN in 15 ml. The pooled fractions were lyophilized and stored at −80 °C. The donor-labeled protein was re-solubilized in the labelling buffer to obtain a protein concentration of approximately 20 μM and the correct protein concentration was determined using the absorbance at 493 nm and the extinction coefficient of the Alexa 488 dye (72000 M^−1^cm^−1^). A 5-fold excess of acceptor dye Alexa 594 C5 maleimide (Invitrogen) was added and samples were incubated for 180 min at (25°C, 300 rpm). Afterwards, the reaction was quenched using β-mercaptoethanol. Again, reversed-phase HPLC was used to remove free label using the same column as described above. The doubly labeled proteins were eluted using with a 25% - 40% ACN gradient. The pooled fractions were lyophilized and stored at −80 °C. The correct labelling of all protein variants was confirmed using ESI mass spectrometry.

### Single-molecule fluorescence spectroscopy

All single-molecule fluorescence experiments were performed with a MicroTime 200 confocal microscope (PicoQuant) equipped with an Olympus IX73 inverted microscope. We either used linearly polarized light from a 485 nm diode laser (LDH-D-C-485, PicoQuant) adjusted to 100 μW (measured at the back aperture of the objective) to excite the donor fluorophore with a repetition rate of 40 MHz or two pulsed lasers controlled by a PDL 828-L “Sepia II” (PicoQuant, Germany) for Pulsed Interleaved Excitation (PIE) experiments^63, 64^. In PIE experiments, the interleaved excitation of the acceptor dye allows us to identify molecules carrying both donor and acceptor dyes. To this end, two light sources, a 485 nm diode laser (LDH-D-C-485, PicoQuant) and a white-light laser (Solea, PicoQuant) set to an excitation wavelength of 595 nm, were used to excite the donor and the acceptor dyes alternatingly at a total repetition rate per period of 20 MHz. The laser intensities were adjusted to 100 μW at 485 nm and 20 μW at 595 nm (Pm100D, Thor Labs). The excitation beam was guided through a major dichroic mirror (ZT 470-491/594 rpc, Chroma) to a 60x, 1.2NA water objective (Olympus) that focuses the beam into the sample. The sample was placed in a homemade cuvette (quartz 25 mm diameter round cover slips, Esco Optics), borosilicate glass 6 mm diameter cloning cylinder (Hilgenberg)) with a volume of 50 μl. Photons emitted from the sample were collected by the same objective and after passing the major dichroic mirror (ZT 470-491/594 rpc, Chroma), the residual excitation light was filtered by a long-pass filter (BLP01-488R, Semrock) and sent through a 100 μm pinhole to remove out-of-focus light. The sample fluorescence was detected with either two channels or four channels for fluorescence anisotropy measurements. Donor and acceptor fluorescence was separated via a dichroic mirror (T585 LPXR, Chroma) and each colour was focused onto a single-photon avalanche diode (SPAD) (Excelitas) with additional bandpass filters: FF03-525/50, (Semrock) for the donor SPAD and FF02-650/100 (Semrock) for the acceptor SPAD. For fluorescence anisotropy measurements, the emission light was first separated into its parallel and perpendicular components with respect to the linearly polarized excitation light via a polarizing beam splitter and each component was separated by two dichroic mirrors into donor and acceptor photons resulting. The arrival time of every detected photon was recorded with a HydraHarp 400 time-correlated single photon counting (TCSPC) module (PicoQuant) and stored with a resolution of 8 ps (16 ps for PIE). The labelled proteins were diluted to a concentration of approximately 50 pM in 20 mM Tris-HCl pH 8.0 (12 mM ionic strength) at the appropriate concentrations of KCl in the absence or presence of unlabelled β-cat. To prevent surface adhesion of the proteins and to maximize photon emission, 0.001% Tween 20 (Pierce) and 20 mM DTT were included in the buffer. Measurements in the presence of β-cat contained in addition 20 mM L-Arg (for a total of 32 mM ionic strength together with the Tris-HCl buffer) to prevent aggregation. All measurements were performed at 23°C.

### Single-molecule identification and FRET corrections

Instrumental imperfections and differences in the brightness of donor and acceptor require a correction of the detected raw photon counts^65, 66^. Although many of the experiments were performed with four detection channels, two donor channels and two acceptor channels with different polarizations, we describe the molecule identification and corrections for a two-channel (channel 1 and 2) setup. However, the extension to four channels is straightforward. These corrections are defined by five parameters: *γ_1_* and *γ_2_* that account for the different detection probabilities of photons from the two dyes, ***β***_*21*_ and ***β***_*12*_, the leakage of donor photons into the acceptor channel 1 and the leakage of acceptor photons into the donor channel 2, respectively, and ***α***, the probability to directly excite the acceptor dye at the wavelength specific for the donor. If *n_1_* and *n_2_* are the detected photons in the acceptor and donor channels, respectively, and *b_1_* and *b_2_* are the background rates in both channels, the corrected photon counts for acceptor and donor (*n_A_*’ and *n_D_*’) are given by

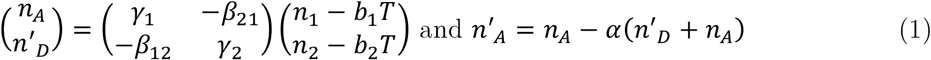

Here, *T* is a time specifying the length of a burst. The correction parameters were determined with two separate samples of the dyes in which their concentrations were adjusted such that both samples have an absorbance of 0.1 at the excitation wavelength (485 nm)^67^. Setting arbitrarily *γ_1_* = 1, we obtain *γ_2_* = 1.12 ± 0.09, ***β***_*21*_ = 0.050 ± 0.003, and ***β***_*12*_ = 0.0021 ± 0.0004 over 5 years with 21 measurements of these correction factors. The probability of directly exciting the acceptor dye at 485 nm is given by ***α*** = ***ϵ_A_*/(*ϵ_A_* + *ϵ_D_*)** where ***ϵ_A_*** and ***ϵ_D_*** are the extinction coefficients of the dyes at the donor excitation wavelength of 485 nm. For our dye pair, ***α*** = 0.05. Transfer efficiency histograms were computed with the fully corrected photon counts according to

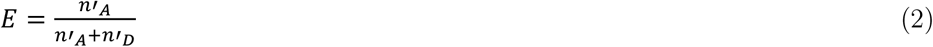

Importantly, the corrections (eq. 1) are already taken into account during burst identification. Bursts were identified from the measured photon traces following Eggeling *et al.*^65^ and Hoffmann *et al.*^66^. If Δ*t_i_* = *t_i_* − *t*_*i-1*_ is the inter-photon time between the *i*^th^ photon and its predecessor, the photon *i* is retained if Δ*t_i_* ≤ *γ*_*j*_(*i*) Δ*t_max_* (Δ*t_max_* = 100 μs) with *γ*_*j*_(*i*) being the correction factor of the *i*^th^ photon detected in channel *j* = [1,2] (see eq. 1). The algorithm then proceeds to the next photon *i* + 1, stops after *n* photons once Δ*t*_*i*+*n*_ > *γ*_*j*_(*i*+*n*) Δ*t_max_*, and provides the total length of the burst by *T* = *t*_*n-1*_ - *t*_*i-1*_. The resulting string of photons is now corrected via eq. 1 using estimated background rates *b_1_* and *b_2_*. The initial guess of *b_1_* and *b_2_* is given by all detected photons in channel 1 and 2, respectively, divided by the total measurement time. A burst is then identified if (*n’_A_* + *n’_D_*) > 100. The photons belonging to this burst are removed from the photon trace and a new guess for *b_1_* and *b_2_* is computed based on the remaining photons. Subsequently, the burst search is performed again with updated background rates. This procedure converges after three iterations to constant background rates and a constant number of identified bursts.

Since the identified bursts also contain molecules for which the acceptor bleached during the transit trough the confocal spot, thus masking the true transfer efficiency, we further cleaned the FRET histograms from these events^68^. For a burst with *n’_D_* donor photons with the arrival times 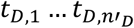 and *n’_A_* acceptor photons with the arrival times 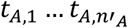, the average arrival times are given by 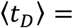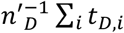 and 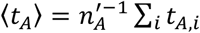. The burst asymmetry is defined by ***α_DA_*** = **〈*t_D_*〉** − **〈*t_A_*〉**. If the acceptor dye bleaches, we clearly find ***α_DA_*** > 0. Taking shot noise into account, the distribution of ***α_DA_*** has a standard deviation given by

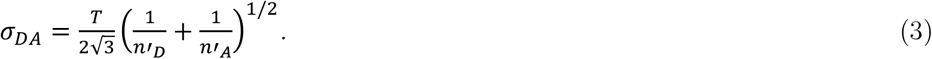

To eliminate molecules with a bleached acceptor, we excluded all molecules for which **|*α_DA_*|** > *σ_DA_*. In addition, only molecules containing active acceptor and donor dyes were included in the analysis. To this end, we computed the donor-acceptor stoichiometry (*S*) for each burst according to

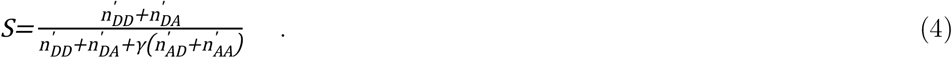

Here, *γ* is a correction factor to account for the different excitation intensities for donor and acceptor. Furthermore, the first subscript indicates the emission and the second subscript indicates the excitation. Only molecules with *S* < 0.8 were used for constructing smFRET histograms.

FRET histograms were fitted with a combination of empirical log-normal and Gaussian distributions^69^. For binding experiments with unlabelled β-cat, the width and position of the FRET-peaks were fixed to minimize the number of free fitting parameters and the area under the fitted histogram curve for each subpopulation was determined using numerical integration. For distance calculations based on the mean transfer efficiencies, the Förster radius measured in water 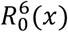 was corrected for the different refractive indices of the solutions according to:

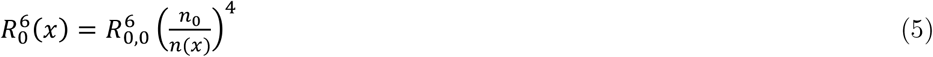

where ***n***(***x***) is the refractive index of the sample at condition ***x***. Refractive indices were measured with an Abbe refractometer (Krüss) and were used to calculate the exact KCl concentrations^70, 71^. The Foerster distance in water is 5.4 nm^72^.

### Two-focus FCS (2fFCS)

2fFCS measurements were conducted on a MicroTime200 (PicoQuant, Berlin) confocal microscope (Olympus). The light of two orthogonally polarized, pulsed diode lasers is combined with a polarization sensitive beam splitter and afterwards coupled into a polarization maintaining single-mode fibre. Both lasers (483 nm, LDH-D-C-485, PicoQuant) are pulsed alternately with a total repetition rate of 40 MHz and a laser power of 30 μW each. Before entering the objective, the laser beam passes through a Nomarski prism (U-DICTHC, Olympus). Afterwards, the two laser beams are focused by the objective thus causing two overlapping excitation volumes with a small lateral shift. Emission light passes through the objective, prism, and the dichroic beamsplitter (see section Single-molecule fluorescence spectroscopy) and is focused onto a pinhole (150 μm). Behind the pinhole the light is collimated and divided by a polarizing beam-splitter and focused onto two single-photon avalanche diodes (SPADs). For each laser focus, the corresponding autocorrelation function and the cross-correlation functions between foci were computed and fitted as described previously^73^. To determine precise diffusion coefficients, we determined the distance between the two foci using the known Stokes radius of the dye Oregon Green and the water viscosity at the known lab-temperature of 23 ± 0.6°C. Fitting of the auto- and cross-correlation functions is performed as described by Dertinger *et al.*^73^. We found a focal distance of 353 nm.

### Determination of the affinity of the core-binding region of E-cad for β-cat

The core-binding region of E-cad with the highest contact probabilities (Fig. 5b) consisted of a 20 amino acid stretch. Since smFRET is not suitable for monitoring conformational changes in such a short peptide, we used 2fFCS to determine the Stokes radius of the peptide at increasing concentrations of β-cat. To minimize that dye-labelling with AlexaFluor 488 massively impacted binding, we used two peptides in which the dye was either place at the N-terminus or at the C-terminus (Supplementary Table 1). In addition, we introduced one serine-glycine repeat between the peptide and the cysteine used for labelling to increase the distance between the dye and the binding competent peptide. The free peptides had an average (average over both labelling constructs) Stokes radius of 1.5 ± 0.2 nm. At the highest β-cat concentration measured in our experiment, the Stokes radius was found to be 1.6 ± 0.2 nm, i.e., unchanged within the error. For comparison, the expected Stokes radius^74^ of β-cat with a molecular weight of 85 kDa is 3.5 nm, which agrees with the Stokes radius of 3.9 nm found for donor-labelled E-cad in complex with β-cat (1 μM). The increased value in presence of E-cad is likely due to the flexible parts of E-cad that render the complex slightly larger than β-cat alone.

### Temperature-controlled smFRET

Temperature-controlled smFRET experiments were conducted in a custom-built system. The design is very similar to a previous version^75, 76^ and includes a temperature controlled sample holder and a second device to control the temperature of the objective. To precisely determine the temperature inside the confocal spot, we measured the diffusion coefficient of the dye Oregon Green (ThermoFisher Scientific) in water using 2f-FCS^73^ (Supplementary Fig. 7). The average of the fits from three independent measurements at each temperature in the range of 0-70 °C were used to determine the viscosity of water given the hydrodynamic radius of Oregon Green (0.6 nm)^77^. The Stokes equation gives a relationship between the hydrodynamic radius ***R***_***S***_, the diffusion coefficient ***D***, and the viscosity of the medium ***η***:

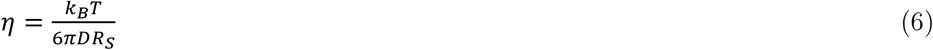

where ***k***_***B***_ is the Boltzmann constant and ***T*** the temperature. On the other hand, the water viscosity ***η*** has a known temperature dependence^78^ given by

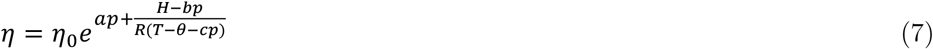

with ***p*** being the pressure in bar, ***T*** the temperature in K, and ***η*_0_**, ***a***, ***b***, ***c***, ***H*** and ***θ*** being constants (***E*** = **4.753 *kJ mol*^−1^**, ***η*_0_** = ***2*.4055 × 10− 5 Pa s**, ***θ*** = **139.*7 K***, ***a*** = **4.42 × 10 ^−4^ bar^−1^**, ***b*** = **9.565 ×10^−4^ *kJ mol*^−1^ bar^−1^**, and ***c*** = **1.24 × 10^−2^ *K bar*^−1^**). When combining eq. 6 and 7, the temperature satisfies an implicit equation that relates the measured diffusion coefficient of Oregon Green to the temperature of the sample, which was used to compute the actual temperature in the confocal spot (Supplementary Fig. 7).

### Determination of average donor-acceptor distances from FRET efficiencies

The mean FRET efficiency of all E-cad variants in the absence of β-cat were converted to distances according to^72^

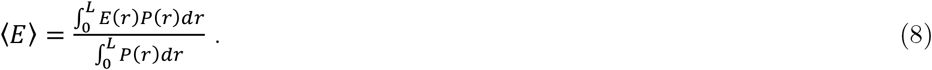

Here, ***E*(*r*)** is the well-known Foerster equation

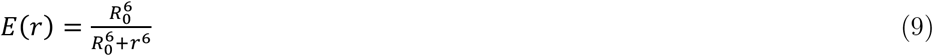

with ***R*_0_** = **5.4 *nm***. The conversion of mean transfer efficiencies to donor-acceptor distances (*R_DA_*) is clearly model-dependent and we determined *R_DA_* using eq. 8 with four polymer models: the Gaussian chain model^72^, the Worm-like chain model^79^, the Sanchez model^25, 80, 81, 82^, and the self-avoiding random walk model (SAW)^79, 83^. The resulting *R_DA_*-distances are all very similar. The model-dependent variation is shown as an error band in Fig. 2d.

### Two-dimensional fluorescence lifetime vs. transfer efficiency plots

To identify whether the observed broadening of the FRET histograms arises from conformational heterogeneity, we computed two-dimensional plots that correlate the donor excited-state lifetime with the measured FRET efficiency^66^. For a fixed distance ***r***, the mean donor lifetime in the presence of acceptor is given by

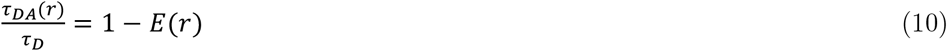

with ***τ_D_*** being the lifetime of the donor in absence of the acceptor. Yet, in presence of a distribution of distances ***P*(*r*)**, the relationship between mean fluorescence lifetime and mean FRET efficiency changes. The mean fluorescence lifetime of the donor in the presence of an acceptor is then given by

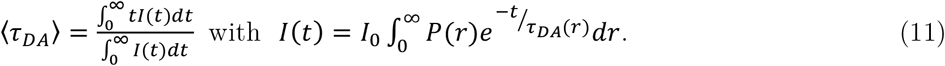

Here, ***I*(*t*)** is the time-dependent fluorescence emission intensity. Simplifying equation (11), the average fluorescence lifetime is given by

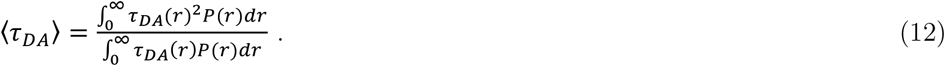

In Fig. 4b, we used the distance distribution for a Gaussian chain given by

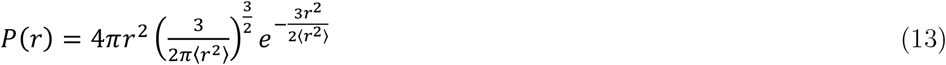

with 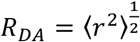 being the average donor-acceptor distance to demonstrate the expected lifetime-FRET correlation for a fully disordered chain.

### Polyampholyte theory

To describe the chain collapse with increasing concentrations of KCl, we fitted the change in donor-acceptor distance of all E-cad variants with a polyampholyte polymer theory developed by Higgs and Joanny^29^

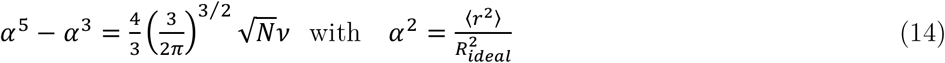

Here, 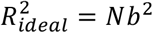 with ***N*** being the number of peptide bonds and ***b* = 0.38 *nm***, being the distance between two C_α_-atoms. Expression (14) measures the deviation of the end-to-end distance with respect to the end-to-end distance of an ideal Gaussian chain. The two-body interaction term (excluded volume) ***ν*** has two contributions ***ν*** = ***ν*_0_ + *ν_el_***, non-electrostatic interactions (***ν*_0_**) and electrostatic interactions (***ν_el_***). The latter is given by Higgs and Joanny as

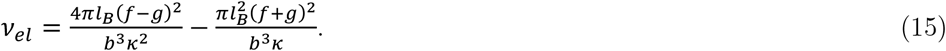

Here, ***f*** and ***g*** are the fractions of positive and negative charges in the sequence of the individual E-cad variants, respectively. The Debye length is defined as 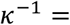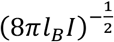, where ***I*** is the ionic strength of the solution and ***l_B_*** is the Bjerrum length, i.e., the distance at which the electrostatic energy of the interaction between two elementary charges equals thermal energy. We use the known relation 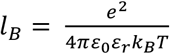, where ***e*** is the elementary charge, ***ε*_0_** is the permittivity of vacuum, ***ε_r_*** is the dielectric constant, ***k_B_*** is Boltzmann’s constant, and ***T*** is the temperature. Hence, eq. 15 is fully determined by the protein sequence and the ionic strength of the medium. To describe the collapse at high concentrations of KCl, we assume a linear dependence between the non-electrostatic two-body interaction term ***ν*_0_ = *c*_1_ + *c*_2_[*I*]** with *I* being the ionic strength of the solution and ***c*_1_** and ***c*_2_** are fitting constants^30^.

### Sequence charge decoration (SCD)

We have used a variant of the sequence charge decoration^34, 35^, *SCD*_lowsalt_,

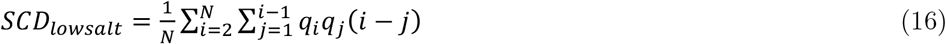

in which *q_i_* and *q_j_* are the charges of the *i*-th and *j*-th amino acids in the sequence. This metric is designed to predict the salt-induced conformational change of the intrinsically disordered proteins due to charge patterning. The chain will compact with addition of salt if *SCD*_lowsalt_ is positive, whereas increasing salt concentration will induce chain expansion if *SCD*_lowsalt_ is negative.

### Nanosecond Fluorescence Correlation Spectroscopy (nsFCS)

We compute subpopulation specific correlation functions. To this end, we are using sample concentrations of around 100-500 pM that still allow us to differentiate populations of molecules that exhibit different FRET efficiencies. In a first step, we identify the photon bursts from individual molecules as described above. For these bursts, the FRET efficiencies are determined and finally the correlation functions for the selected subset of molecules were computed. By distributing the photons onto two donor and two acceptor detectors, dead time and after-pulsing of the SPADs are avoided. The cross-correlation functions were fitted using

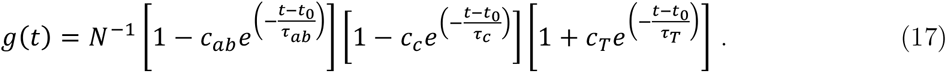

The three terms in brackets describe photon antibunching (***ab***), conformational dynamics (***c***), and triplet blinking of the dyes (***T***). In addition, *N* is the effective number of molecules in the confocal volume and ***c_ab_***, ***c_c_***, and ***c_T_*** are amplitudes. The correlation times ***τ_ab_***, ***τ_c_***, ***τ_T_*** and the time origin ***t*_0_** are fitting parameters. To graphically compare different experiments, the correlation functions in Fig. 4a were normalised by *N* and the triplet term.

### Determining the intra-chain diffusion coefficient of E-cad ABC from nsFCS

We determined the intra-chain diffusion coefficient of free E-cad ABC by modelling its dynamics as a diffusive process in the potential of mean force given by the donor-acceptor distance distribution that resulted as a best fit from the CG-simulations (Fig. 6c). In this picture, the measured correlation time ***τ_c_*** obeys a direct relationship to the intra-chain diffusion coefficient ***D*** given by^84^

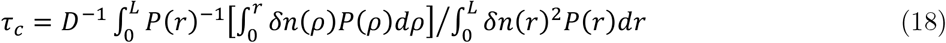

with

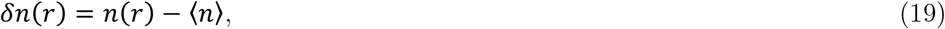

where ***P*(*r*)** is the donor-acceptor distance distribution determined from the CG-simulation and ***n*(*r*)** is the donor photon rate. The latter is determined from the photophysics of the FRET system with donor (D) and acceptor (A) that includes the four states DA, D*A, D*A*, DA* where the asterisk indicates an excited state. The rate matrix for this system is given by (in the basis given above)

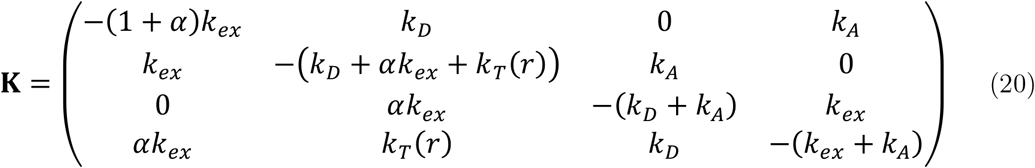

where ***k_ex_* = 0.018 *ns*^−1^** is the donor excitation rate, ***k_D_* = 0.28 *ns*^−1^** is the donor emission rate, ***k_A_* = 0.25 *ns*^−1^** is the acceptor emission rate, and ***α* = 0.05** is the direct excitation probability. The energy transfer rate is given by ***k_T_*(*r*) = *k_D_*(*R*_0_/*r*)^6^**. The steady-state vector of the four states **p_ss_** is the solution of the algebraic system 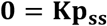 where **0** is the null vector. The photon rate ***n*(*r*)** of the system is then given by 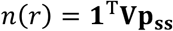 where

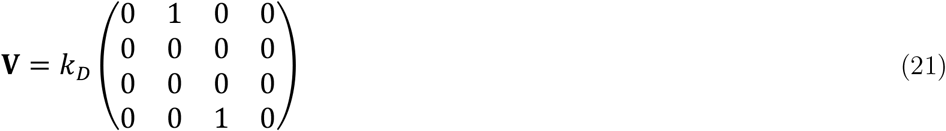

is the donor photon detection matrix. We determined the intra-chain diffusion coefficient ***D*** of free E-cad ABC (Fig. 6c) by numerically solving eq. 18.

### Recurrence analysis of single particles (RASP)

The recurrence analysis of single particles (RASP) uses the fact that a freely diffusing molecule can be observed multiple times in a single-molecule experiment. Once a molecule leaves the observation volume, the chance of it returning to the confocal spot is greater than the chance of detecting a new molecule for short time intervals (Supplementary Fig. 4a-b). To extract kinetics based on this effect, we binned our data in 100 μs bins and computed the bin-time autocorrelation function ***G***(***τ***) that contains the information of the likelihood to observe two molecules separated by the delay time ***τ***

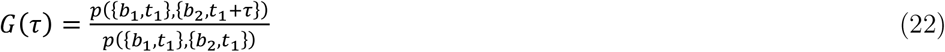

Here, ***p*({*b*_1_, *t*_1_}, {*b*_2_, *t*_1_ + *τ*})** denotes the joint probability of observing two bins ***b*_1_** and ***b*_2_** at times ***t*_1_** and ***t*_1_ + *τ***, respectively, and ***p*({*b*_1_, *t*_1_})** and ***p*({*b*_2_, *t*_1_})** are the probabilities of detecting ***b*_1_** and ***b*_2_**, respectively, at time ***t*_1_**. This correlation function can be converted to ***P***_*same*_(*τ*), i.e., the probability that bins separated by the time *τ* originate from the same molecule^42^ (Supplementary Fig. 4b):

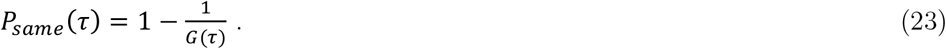

During the time between the first and the second detection, the molecule can change its conformation, thus causing a different FRET efficiency in the second detection. To quantify the conformational dynamics with RASP, we define two regions of the broad FRET histograms of the E-cad/β-cat complex, region 1 and region 2 (Supplementary Fig. 4c). Bins originating from the FRET efficiency chosen in region 1 are selected (E0-bins) and recurrence histograms are calculated from the photons in bins (E1-bins) at a time delay ! after the initial bins (Supplementary Fig. 4d). The recurrence FRET-histograms were globally fitted with a sum of a log-normal function for molecules that lack an active acceptor dye (E = −0.05) and two Gaussian functions for the populations from regions 1 and 2. The positions, widths, and asymmetries were determined from the histograms of both regions at the initial time window, i.e., at *τ* = 0. The relative populations of the molecules in region 1 and region 2 were obtained by integrating the respective sub-populations. Supplementary Fig. 4e depicts the time course of the fraction of molecules in region 1 after originally selecting molecules in region 2 (red) or molecules in region 1 (blue), respectively. By increasing the time delay !, the fraction of molecules in region 1 when initially selecting region 1 decrease and those in region 1 when selecting region 2 increase. The observed kinetics ***P_m_*(*t*)** is a combination of two contributions: conformational switching between the two selected regions, characterized by ***P_conf_*(*t*)**, and the time-dependent likelihood that two observed bins or “snapshot” arise from the same molecule, ***P_same_*(*t*)**. Since we can compute ***P_same_*(*t*)** directly from the burst time autocorrelation function of our data, we can determine ***P_conf_*(*t*)**, the actual kinetics of conformational switching. Thus, the observed increase in the population of molecules in region 1 in the recurrence histograms after initially selecting molecules in region 2 is given by

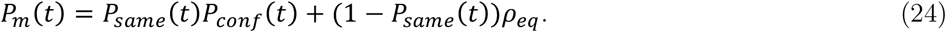

Here, ***P_m_*(*t*)** is the measured time decay, ***P_same_*(*t*)** is the probability that the same molecule recurs, ***P_conf_*(*t*)** is the kinetics of switching to region 1 due to conformational dynamics, 1 − ***P_same_*(*t*)** is the probability that a new molecule enters the confocal volume, and *ρ_eq_* is the probability that a new molecule is in region 1, which is given by the equilibrium probability that a molecule is in region 1. For a better visualization and easier analysis, we directly determine the ‘true’ kinetics of switching ***P_conf_*(*t*)** that we obtain by re-arranging equation (24) for ***P_conf_*(*t*)**. In addition, we average the two kinetic traces, i.e., the increase of molecules in region 1 after initial selection of molecules from region 2 and the depletion of molecules in region 1 after initial selection of molecules from region 1. The resulting averaged traces are shown in Fig. 4c.

### Fitting RASP kinetics by numerically solving the Smoluchowski equation

As an alternative to exponential fitting, we also fitted the RASP kinetics of the variant E-cad ABC bound to β-cat by solving the Smoluchowski equation

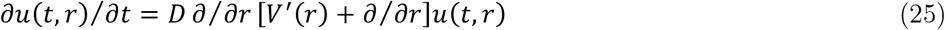

with ***u*(*t, r*)** being the time-dependent donor-acceptor distance distribution and ***V*(*r*)** = − ln ***P*(*r*)** being the potential of mean force with ***P*(*r*)** being the normalized equilibrium donor-acceptor distance distribution obtained as a best fit from the CG-model (Fig. 6b inset). Equation 25 was solved with the boundary conditions

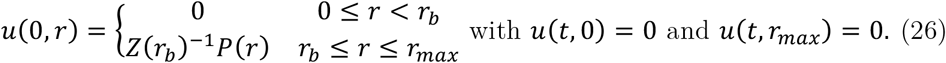

Here, ***Z*(*r_b_*)** is a normalization factor, ***r_b_*** defines the distance that separates high-FRET (region 2) (***r* < *r_b_*)** from low-FRET (region 1) (***r* ≥ *r_b_***) molecules, and ***r_max_*** is the upper integration limit (20 nm). An analytical expression for ***P*(*r*)** suitable to efficiently solve eq. 25 was obtained by fitting the distance distribution for E-cad ABC from the CG-model with the superposition of three Gaussian functions. The fraction of low-FRET molecules ***f*(*t*)**, i.e., molecules from region 1, was obtained from

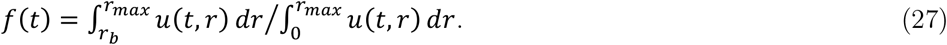

The model contains two free parameters, the diffusion coefficient ***D*** and the separation distance ***r_b_***. The latter quantity was determined prior to fitting by numerically solving

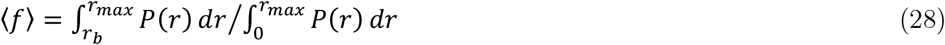

for ***r_b_*** where **〈*f*〉** is the average fraction of the last 2 ms of the kinetic time traces, i.e., from 8 – 10 ms. The resulting value for ***r_b_*** was then used for fitting. We performed an error estimation of the diffusion coefficient using bootstrapping the kinetic time traces with *n* = 3.

### Molecular simulations with the CG-model

We parameterized a coarse-grained (CG) model to characterize the interactions between E-cad and β-cat. The model was based on our recently developed HPS model^85^ in which each amino acid was represented as a bead. The potential energy can be written as,

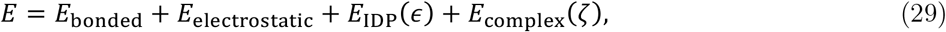

in which there are three types of interactions for disordered E-cad: bonded, electrostatic and short-range pairwise interactions, and one additional term for the interactions between E-cad and β-cat. The bonded interactions were modelled by a harmonic potential with a bond length of 3.8 Å and a spring constant of 10 kJ/Å^2^. The electrostatic interactions were modelled with a Coulombic term using Debye-Hückel electrostatic screening^86^. The short-range pairwise interactions were modelled using Ashbaugh-Hatch functional form^87^ and the depths of the potential minimums are proportional to the amino acid hydrophobicity values (***λ***)^88^. The free parameter ***ε***, which was originally tuned according to the size of the IDPs, has been adjusted to 0.16 kcal/mol using the experimental FRET efficiencies of the six E-cad segments (Supplementary Fig. 8). To simulate β-cat in the CG model, we represented the structured part of β-cat in the experimentally solved structure (PDB: 1i7x)^23^ as a rigid body in the simulation. The interactions between E-cad and β-cat were also modelled based on the native interactions in the same structure. A native interaction was assumed if the Cα distance between two amino acids is smaller than 1.2 times the sum of the amino acid radii used in the CG model. For every pair of amino acids that form native interactions in the complex, we have introduced one rigid and one flexible CG-models, with different types of potential energy functions. First for the rigid model, a harmonic potential energy function with the native interaction distance as the minimum position and a large spring constant of 20 kcal/mol were introduced. This effectively rigidified all the native contacts as seen in both X-ray structures. Second for the flexible model, a 12-6 Leonard-Jones (LJ) potential energy function with the native interaction distance as the minimum position was introduced. The depth of the LJ-well, i.e., the interaction strength ζ was adjusted to 0.6 kcal/mol according to the FRET efficiencies of the six segments of E-cad in the complex at an ionic strength of 82 mM. The deviations to the experimental FRET efficiencies for both models were shown in Supplementary Fig. 8. The CG simulation of E-cad alone was run for 10 μs and the complex simulation was run for 20 μs, using LAMMPs^89^ and HOOMD-Blue^90^ to benefit from both CPU and GPU resources. The first 1 μs of the simulation was always excluded from the analysis. To obtain the FRET efficiencies from the simulations, the linker of both dyes were taken to be equivalent 9 amino acids^91^, i.e., 4.5 amino acids per linker.

### Rouse model with surface interactions

Here we show that for a simple bead-and-spring model, contacts with a surface increase the effective spring constant. In one dimension, the Langevin equation for the coordinate of the *k*^th^ bead (***x_k_* = *x_k_*(*t*)**) that contacts its neighbour beads is well known^46, 92^. All beads also contact a specific location on a surface (***y* = 0**), which, for simplicity, is identical for all beads and does not fluctuate in time

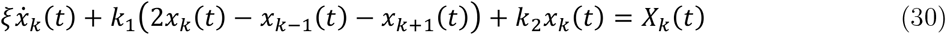

Here, *ξ* is a friction coefficient and *k_1_* and *k_2_* are spring constants for bonds in the chain and with the surface, respectively. ***X_k_*** is Gaussian distributed white noise. Assuming a circular chain^46^, taking the Fourier transform in bead number (***k* → *q***) and time (***t* → *ω***), and rearranging, we obtain

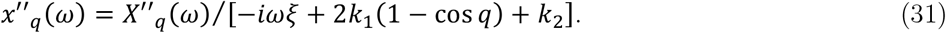

The relaxation time for the largest modes of such a chain is then given by ***τ*_*q*_ = *ξ/k_eff_*** with the effective spring constant ***k_eff_* = *k*_1_*q*^2^ + *k*_2_**, showing that the spring constant of a chain is affected by surface interactions.

## Data availability

Data supporting the findings of this study are available from the corresponding authors upon reasonable request. A reporting summary of this article is available as a Supplementary information file. Source data are provided with this paper.

## Code availability

A custom WSTP add-on for Mathematica (Wolfram Research) used for the analysis of single-molecule fluorescence data is available at https://schuler.bioc.uzh.ch/programs/. The implementation of the coarse-grained HPS model in HOOMD-Blue can be downloaded at https://bitbucket.org/jeetain/hoomd_slab_builder.

## Acknowledgements

We thank William I. Weis for providing the E-cad plasmid and Dirk Görlich for the plasmid containing the SUMO protease. We are also grateful to Nir London and Christian Dubiella for their help with mass spectrometry. In addition, we enjoyed critical discussions with Gilad Haran, Amnon Horovitz, Koby Levy, Benjamin Schuler, Robert Best, and Philipp Selenko. This work was supported by the Israel Science Foundation (grant no. 1549/15), the Benoziyo Fund for the Advancement of Science, the Carolito Foundation, the Leir Charitable Foundation, and the Koshland family. W.Z. acknowledges the support from the National Science Foundation (MCB-2015030) and the research computing at Arizona State University for providing HPC.

---

**Supplementary Fig. 1:**
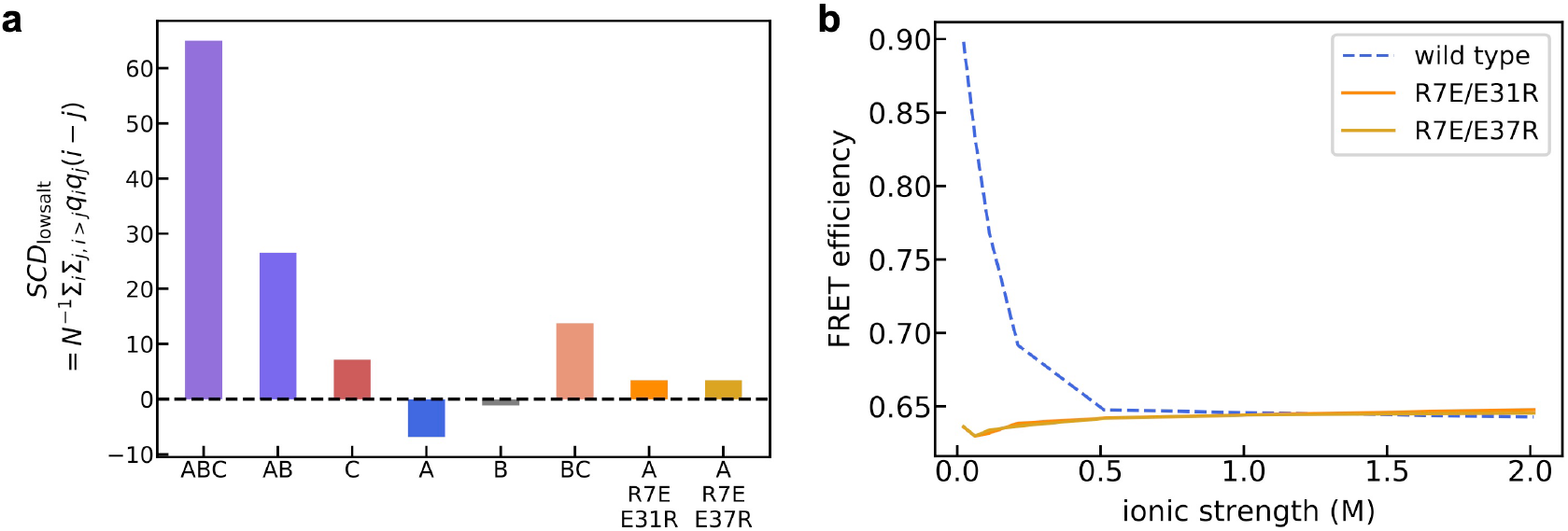
Sequence charge decoration metric (SCD_lowsalt_, see Methods) for all E-cad segments computed based on the amino acid sequence (**a**) and salt-dependent FRET efficiencies of wild type E-cad and two charge-swapped variants of the A-segment (**b**).

**Supplementary Fig. 2.**
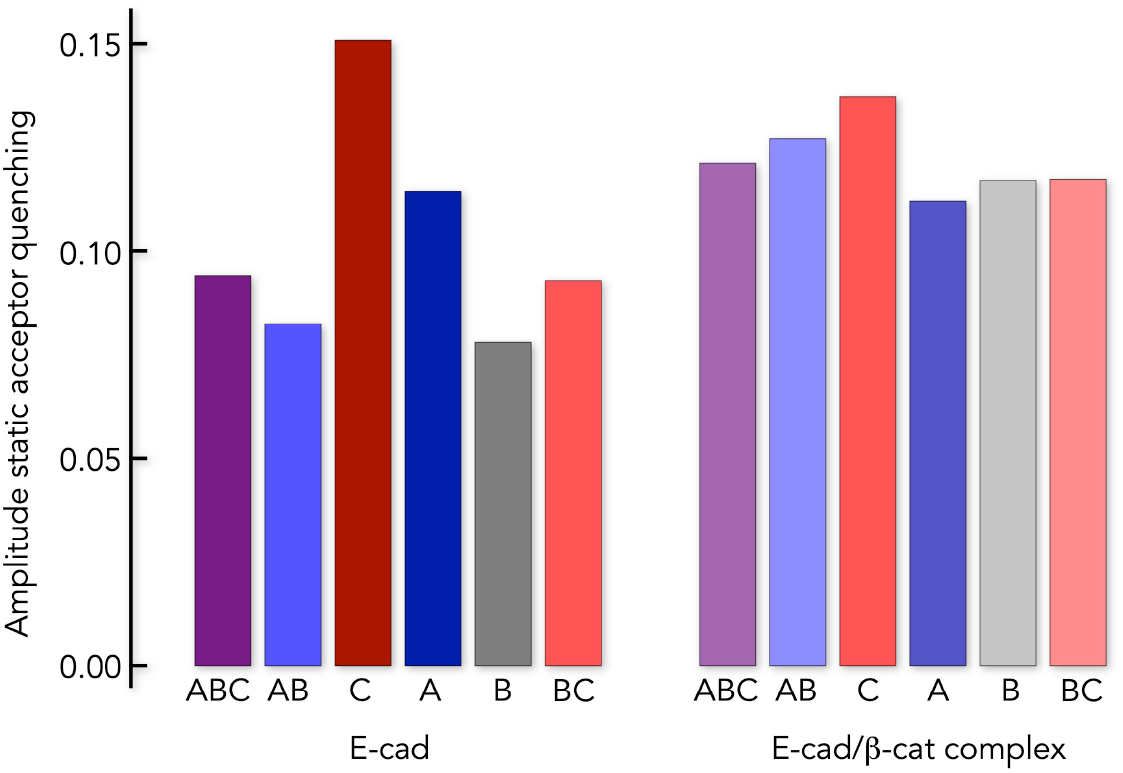
nsFCS of all E-cad variants in the absence (left) and presence (right) of β-cat after exciting the acceptor directly. Quantitative comparison of the quenching amplitudes show that the C-segment has the highest amplitude in free E-cad. All quenching amplitudes are elevated in complex with β-cat.

**Supplementary Fig. 3.**
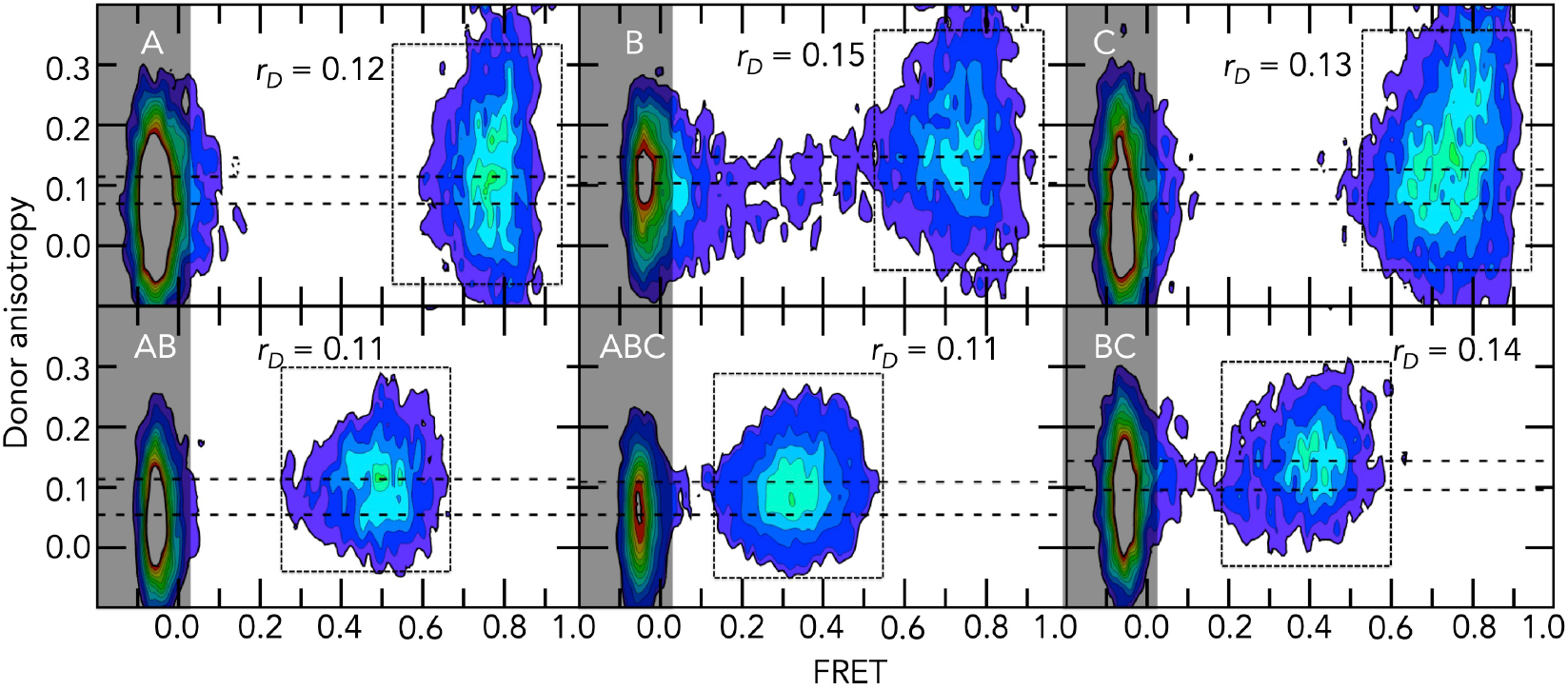
Steady-state donor anisotropy of the six E-cad variants (indicated) in complex with β-cat. The population at a FRET efficiency close to zero arises from molecules that lack an active acceptor (gray shaded area). The second population at higher FRET efficiency (box) results from E-cad molecules with donor and acceptor in complex with β-cat. For all variants, the steady-state anisotropies are between 0.1 and 0.2, indicating no substantial restriction in the rotational freedom of the donor dye. The average donor anisotropies of the FRET-population is indicated.

**Supplementary Fig. 4.**
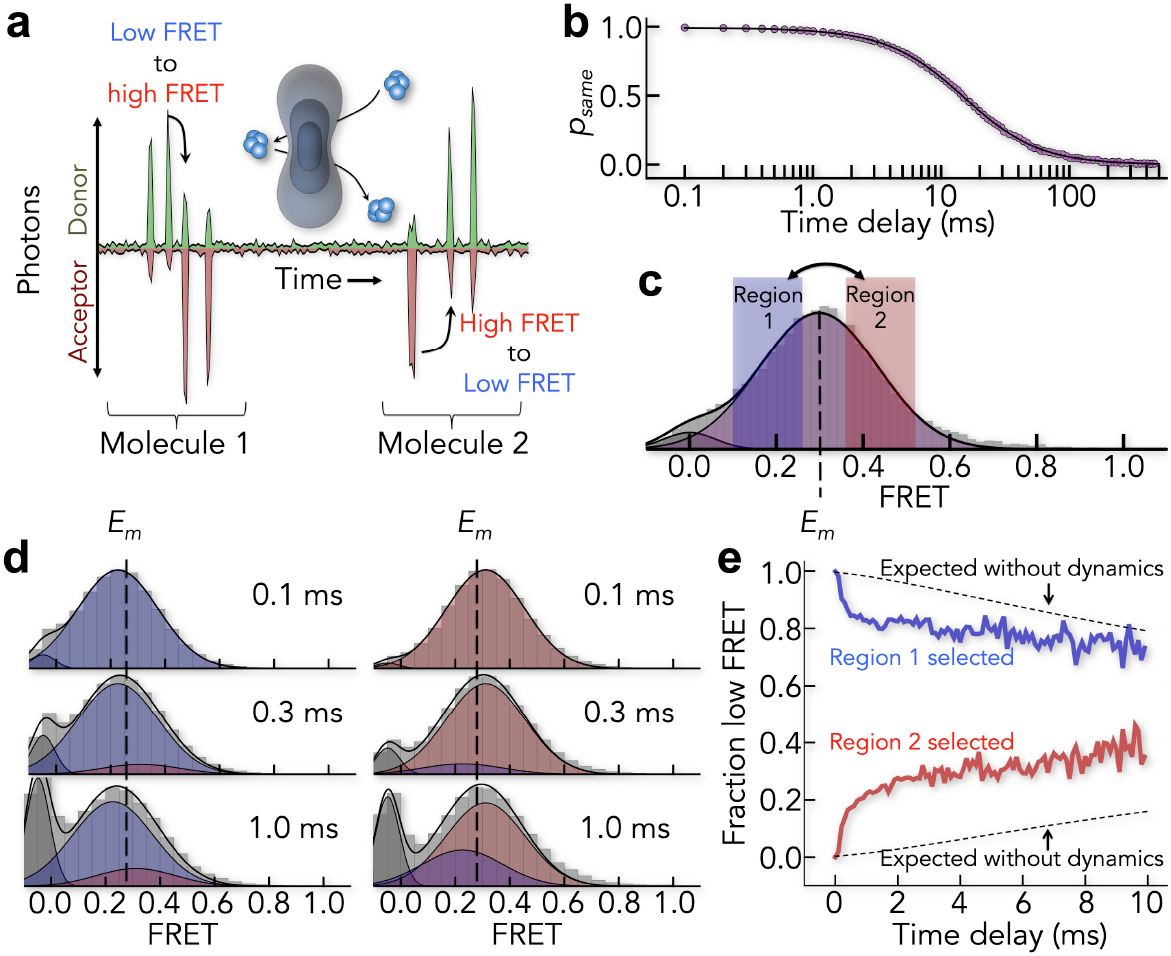
Recurrence analysis of single particles (RASP) of the E-cad/β-cat complex. (**a**) Scheme of single molecules that are detected while they diffuse through the confocal volume. After the initial molecule exits the confocal volume, it can either return or a new molecule can enter. (**b**) Time-dependent probability to detect the same molecule several time decays. (**c**) Equilibrium FRET histogram of the E-cad/β-cat complex. For RASP, molecules were selected either from region 1 (low FRET) or from region 2 (high FRET). The dashed line indicates the mean FRET position (*E_m_*). (**d**) Time-dependent recurrence histograms of molecules selected from region 1 (left) and region 2 (right). Solid lines are fits with a superposition of Gaussian peaks. The dashed line indicates the mean FRET position (*E_m_*) of the equilibrium FRET histogram shown in c. (**e**) Time decays of the fraction of molecules in region 1 (low-FRET) after initially selecting molecules from region 1 (blue) or region 2 (red). The dashed line indicates the expected kinetics in the absence of conformation dynamics as computed from the *p_same_* (see panel b).

**Supplementary Fig. 5.**
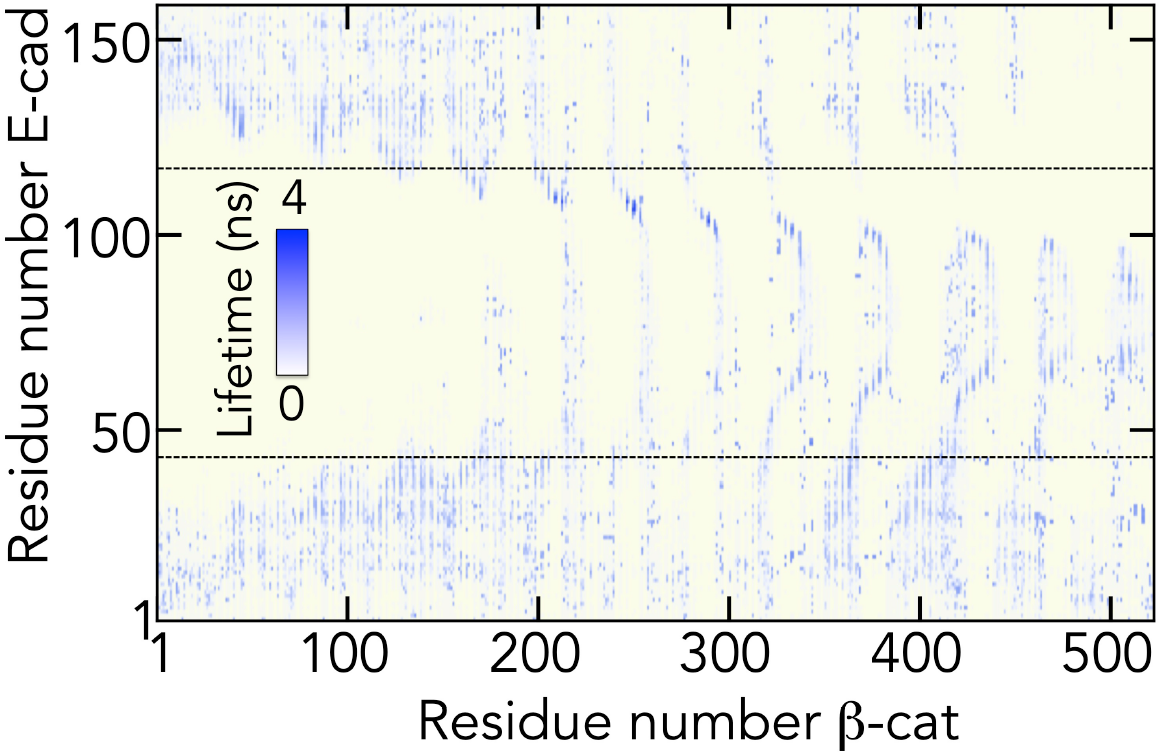
Contact lifetimes in the E-cad/β-cat complex from the flexible CG-model. The contact lifetime map shows a significant distribution of contacts across the whole β-cat surface.

**Supplementary Fig. 6.**
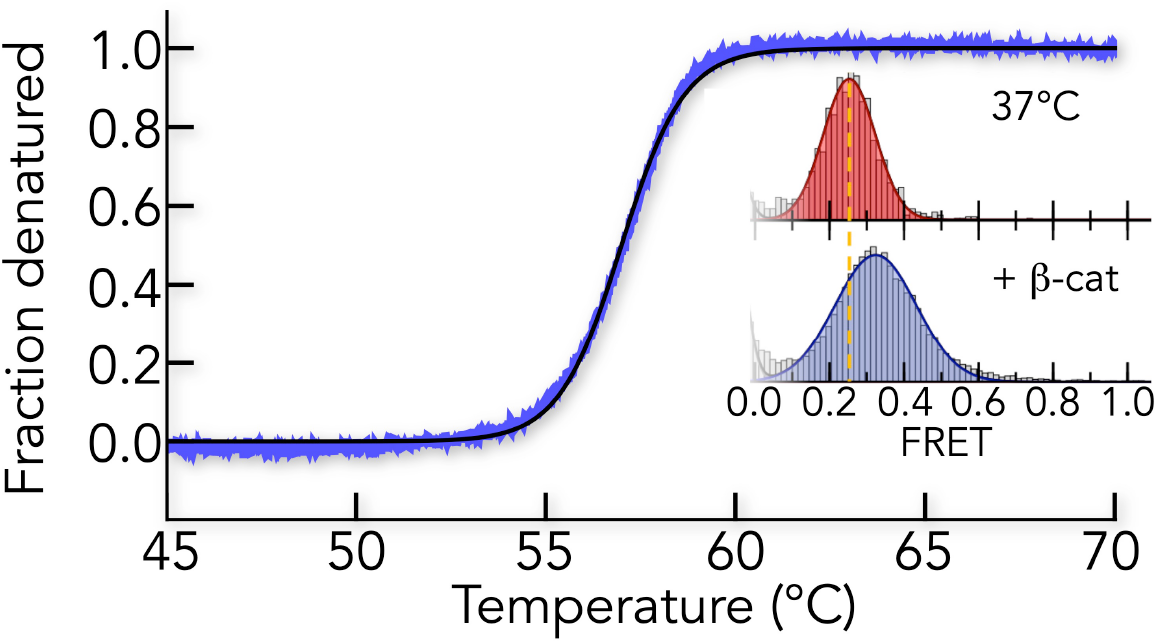
Thermal stability of β-cat monitored with nano-differential scanning fluorimetry (nanoDSF). The melting curve shows a denaturation at 57°C, indicating that β-cat is stable at the highest temperature used in the temperature-dependent RASP experiments (main text Fig. 6). Inset: SmFRET histograms of E-cad ABC at 37°C in the absence (blue) and presence (red) of β-cat (200 nM). The significant broadening of the FRET-distribution at 37°C and the shift to higher FRET-values indicate binding of E-cad at 37°C. Solid lines are fits with a superposition of two Gaussian peaks.

**Supplementary Fig. 7.**
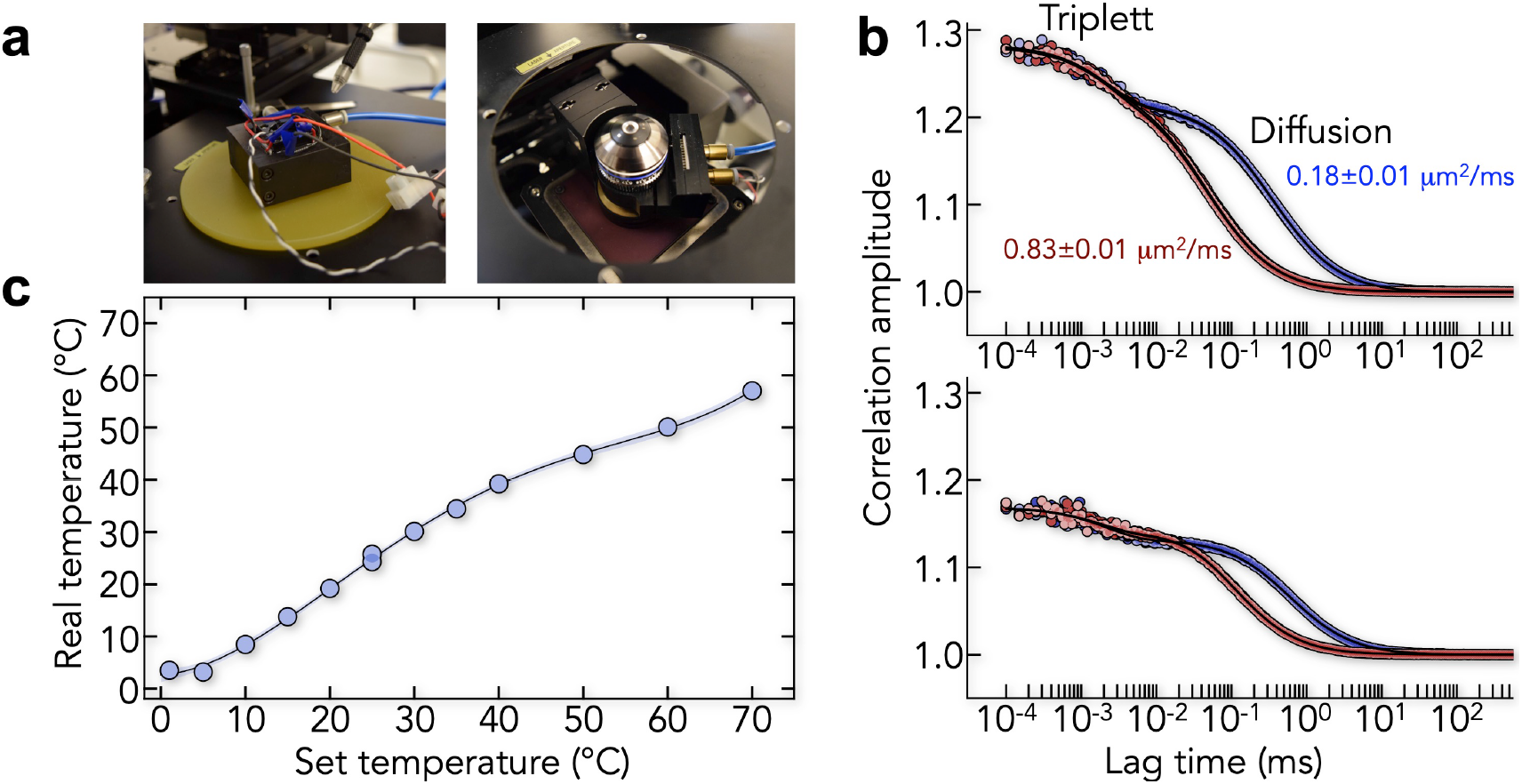
Calibration of smFRET experiments at different temperatures. (**a**) Peltier-controlled sample holder (left) and device to control the temperature of the objective (right). (**b**) Dual-focus FCS autocorrelation functions (top) and cross-correlation functions (bottom) of the dye Oregon Green at set temperatures 0°C (blue circles) and 70°C (red circles). Solid lines are fits as described in the Methods section. The diffusion coefficients at the two temperatures are indicated. (**c**) Temperature calibration curve obtained by determining the diffusion coefficient of Oregon Green using 2fFCS. The x-axis shows the temperature set on the sample controller. The objective controller is set to the same temperature up to a set-temperature of 40°C. For higher temperatures, the objective is kept at a set-temperature of 40°C. The y-axis shows the temperature determined based on the diffusion coefficient of Oregon Green and the known temperature dependence of the water viscosity (see Methods). The solid line is a polynomial fit of fourth order to the data and the blue shaded are indicates the 90% confidence band.

**Supplementary Fig. 8.**
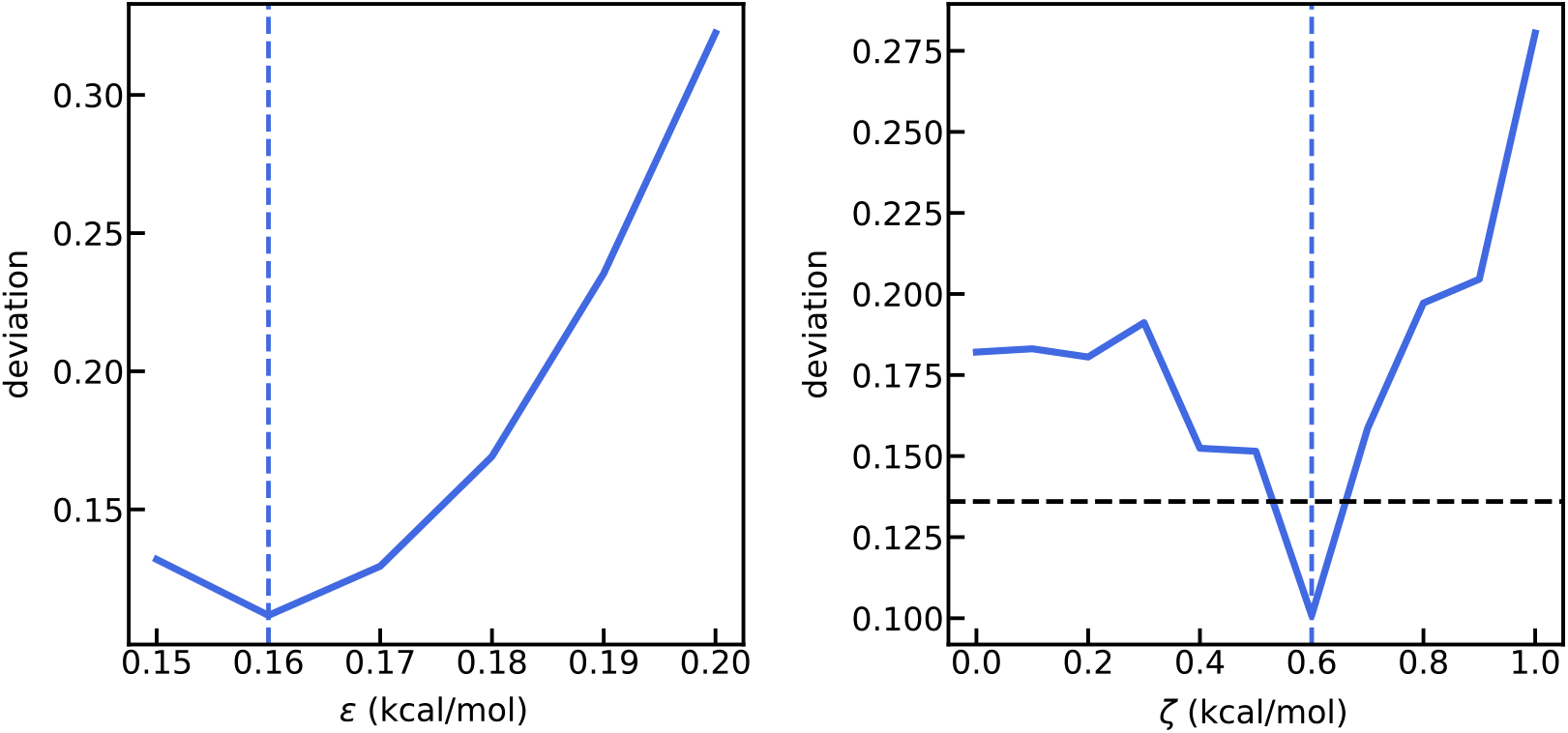
Parameterization of the coarse-grained (CG) model. Left: Deviation of the calculated FRET-values from the experimental data of the six constructs when varying the interaction strength (*ε*) of E-cad only. Right: The deviation of the model from the FRET experiment of the six constructs at the experimental ionic strength of 82 mM when varying the interaction strength (ζ) between E-cad and β-cat for the flexible CG-model (see Methods). The blue dashed lines indicate the minimum and the black dashed lines indicate the deviation for the rigid CG-model.

**Supplementary Table 1.**
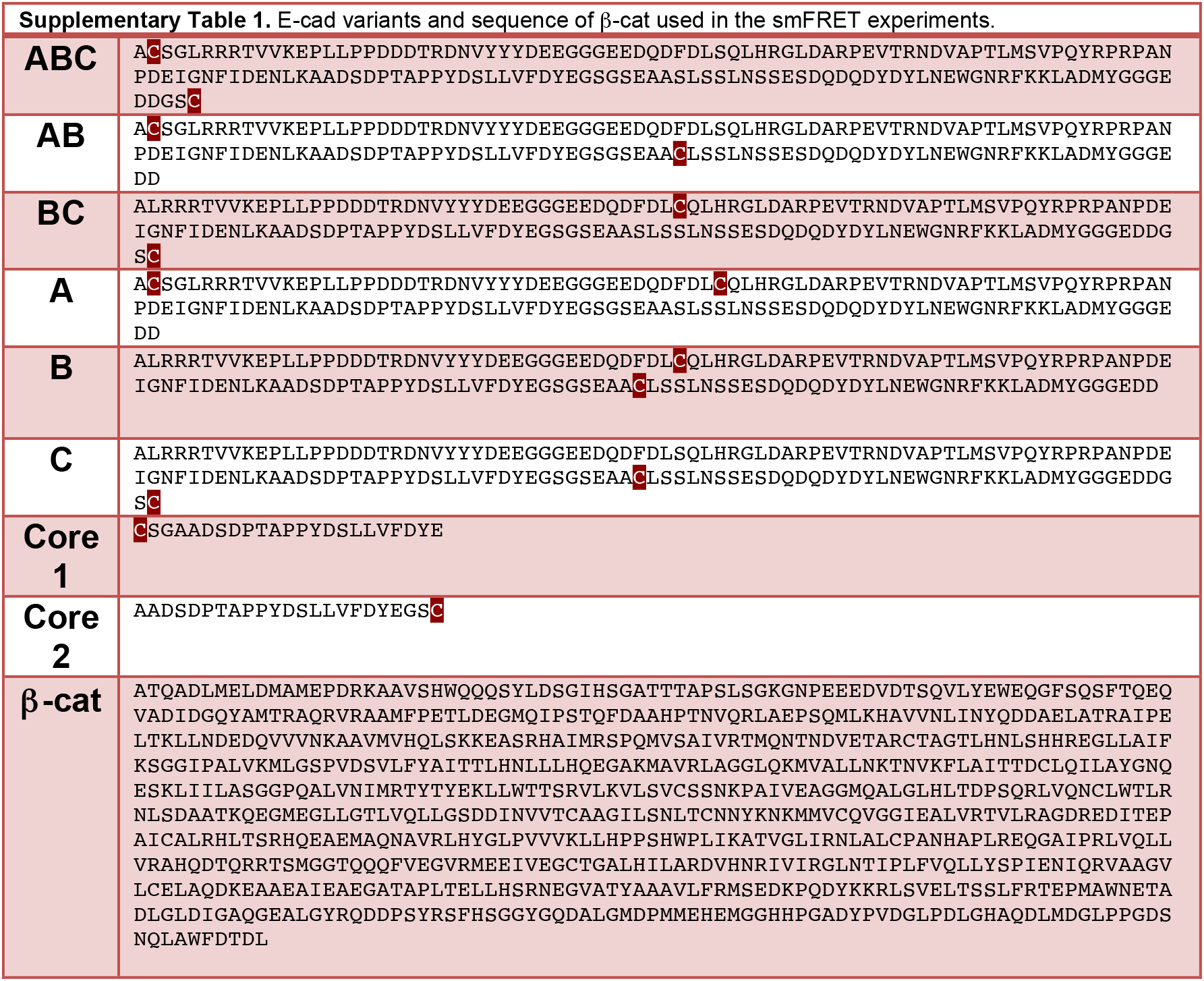
E-cad variants and sequence of β-cat used in the smFRET experiments.

## Notes

### Competing Interest Statement

The authors have declared no competing interest.

## References

1. Schuler B, et al. Binding without folding - the biomolecular function of disordered polyelectrolyte complexes. Curr Opin Struct Biol 60, 66–76 (2020).

2. van der Lee R, et al. Classification of Intrinsically Disordered Regions and Proteins. Chem Rev 114, 6589–6631 (2014).

3. Csizmok V, Follis AV, Kriwacki RW, Forman-Kay JD. Dynamic Protein Interaction Networks and New Structural Paradigms in Signaling. Chem Rev 116, 6424–6462 (2016).

4. Levy Y, Onuchic JN, Wolynes PG. Fly-casting in protein-DNA binding: frustration between protein folding and electrostatics facilitates target recognition. J Am Chem Soc 129, 738–739 (2007).

5. Borgia A, et al. Extreme disorder in an ultrahigh-affinity protein complex. Nature 555, 61–66 (2018).

6. Holmstrom ED, Liu Z, Nettels D, Best RB, Schuler B. Disordered RNA chaperones can enhance nucleic acid folding via local charge screening. Nat Comms 10, 2453–2411 (2019).

7. Mittag T, et al. Dynamic equilibrium engagement of a polyvalent ligand with a single-site receptor. Proc Natl Acad Sci USA 105, 17772–17777 (2008).

8. Hendus-Altenburger R, et al. The human Na + /H + exchanger 1 is a membrane scaffold protein for extracellular signal-regulated kinase 2. BMC Biol 14, 1–17 (2016).

9. Milles S, et al. Plasticity of an ultrafast interaction between nucleoporins and nuclear transport receptors. Cell 163, 734–745 (2015).

10. Nusse R, Clevers H. Wnt/β-Catenin Signaling, Disease, and Emerging Therapeutic Modalities. Cell 169, 985–999 (2017).

11. McCrea PD, Turck CW, Gumbiner B. A homolog of the armadillo protein in Drosophila (plakoglobin) associated with E-cadherin. Science 254, 1359–1361 (1991).

12. Kemler R. From cadherins to catenins: cytoplasmic protein interactions and regulation of cell adhesion. Trends Genet 9, 317–321 (1993).

13. Behrens J, et al. Functional interaction of beta-catenin with the transcription factor LEF-1. Nature 382, 638–642 (1996).

14. Aberle H, Butz S, Stappert J, Weissig H, Kemler R, Hoschuetzky H. Assembly of the cadherin-catenin complex in vitro with recombinant proteins. J Cell Sci 107, 3655–3663 (1994).

15. Ozawa M, Baribault H, Kemler R. The cytoplasmic domain of the cell adhesion molecule uvomorulin associates with three independent proteins structurally related in different species. EMBO J 8, 1711–1717 (1989).

16. Rimm DL, Koslov ER, Kebriaei P, Cianci CD, Morrow JS. Alpha 1(E)-catenin is an actin-binding and -bundling protein mediating the attachment of F-actin to the membrane adhesion complex. Proc Natl Acad Sci USA 92, 8813–8817 (1995).

17. Huber AH, Nelson WJ, Weis WI. Three-dimensional structure of the armadillo repeat region of beta-catenin. Cell 90, 871–882 (1997).

18. Behrens J, et al. Functional interaction of beta-catenin with the transcription factor LEF-1. Nature 382, 638–642 (1996).

19. Hoschuetzky H, Aberle H, Kemler R. Beta-catenin mediates the interaction of the cadherin-catenin complex with epidermal growth factor receptor. J Cell Biol 127, 1375–1380 (1994).

20. Hülsken J, Birchmeier W, Behrens J. E-cadherin and APC compete for the interaction with beta-catenin and the cytoskeleton. J Cell Biol 127, 2061–2069 (1994).

21. Aberle H, Schwartz H, Hoschuetzky H, Kemler R. Single amino acid substitutions in proteins of the armadillo gene family abolish their binding to alpha-catenin. J Biol Chem 271, 1520–1526 (1996).

22. MacDonald BT, Tamai K, He X. Wnt/β-Catenin Signaling: Components, Mechanisms, and Diseases. Dev Cell 17, 9–26 (2009).

23. Huber AH, Weis WI. The structure of the beta-catenin/E-cadherin complex and the molecular basis of diverse ligand recognition by beta-catenin. Cell 105, 391–402 (2001).

24. DeForte S, Uversky VN. Resolving the ambiguity: Making sense of intrinsic disorder when PDB structures disagree. Protein Sci 25, 676–688 (2016).

25. Hofmann H, Soranno A, Borgia A, Gast K, Nettels D, Schuler B. Polymer scaling laws of unfolded and intrinsically disordered proteins quantified with single-molecule spectroscopy. Proc Natl Acad Sci USA 109, 16155–16160 (2012).

26. Ha, Thirumalai. Conformations of a polyelectrolyte chain. Phys Rev A 46, R3012–R3015 (1992).

27. Hofmann H, Golbik R, Ott M, Hübner C, Ulbrich-Hofmann R. Coulomb forces control the density of the collapsed unfolded state of barstar. J Mol Biol 376, 597–605 (2008).

28. Müller-Späth S, et al. Charge Interactions can Dominate the Dimensions of Intrinsically Disordered Proteins. Proc Natl Acad Sci USA 107, 14609–14614 (2010).

29. Higgs PG, Joanny J-F. Theory of polyampholyte solutions. J Chem Phys 94, 1543–1554 (1991).

30. Vancraenenbroeck R, Harel YS, Zheng W, Hofmann H. Polymer effects modulate binding affinities in disordered proteins. Proc Natl Acad Sci USA 116, 19506–19512 (2019).

31. Bhattacharjee A, Kundu P, Dua A. Self-consisten theory of structures and transitions in weak polyampholytes. Macromol Theory Simul 20, 75–84 (2011).

32. Samanta HS, Chakraborty D, Thirumalai D. Charge fluctuation effects on the shape of flexible polyampholytes with applications to intrinsically disordered proteins. J Chem Phys 149, 163323 (2018).

33. Das RK, Pappu RV. Conformations of intrinsically disordered proteins are influenced by linear sequence distributions of oppositely charged residues. Proc Natl Acad Sci U S A 110, 13392–13397 (2013).

34. Sawle L, Ghosh K. A theoretical method to compute sequence dependent configurational properties in charged polymers and proteins. J Chem Phys 143, 085101 (2015).

35. Huihui J, Firman T, Ghosh K. Modulating charge patterning and ionic strength as a strategy to induce conformational changes in intrinsically disordered proteins. J Chem Phys 149, 085101 (2018).

36. Gomes G-NW, et al. Conformational Ensembles of an Intrinsically Disordered Protein Consistent with NMR, SAXS, and Single-Molecule FRET. J Am Chem Soc 142, (2020).

37. Choi H-J, Huber AH, Weis WI. Thermodynamics of β-Catenin-Ligand Interactions THE ROLES OF THE N- AND C-TERMINAL TAILS IN MODULATING BINDING AFFINITY. J Biol Chem 281, 1027–1038 (2006).

38. Nettels D, Gopich I, Hoffmann A, Schuler B. Ultrafast dynamics of protein collapse from single-molecule photon statistics. Proc Natl Acad Sci USA 104, 2655–2660 (2007).

39. Schuler B, Soranno A, Hofmann H, Nettels D. Single-Molecule FRET Spectroscopy and the Polymer Physics of Unfolded and Intrinsically Disordered Proteins. Annu Rev Biophys 45, 207–231 (2016).

40. Haenni D, Zosel F, Reymond L, Nettels D, Schuler B. Intramolecular Distances and Dynamics from the Combined Photon Statistics of Single-Molecule FRET and Photoinduced Electron Transfer. J Phys Chem B 117, 13015–13028 (2013).

41. Hillger F, et al. Probing protein-chaperone interactions with single-molecule fluorescence spectroscopy. Angew Chem Int Ed Engl 47, 6184–6188 (2008).

42. Hoffmann A, et al. Quantifying heterogeneity and conformational dynamics from single molecule FRET of diffusing molecules: recurrence analysis of single particles (RASP). Phys Chem Chem Phys 13, 1857–1871 (2011).

43. Grossman-Haham I, Rosenblum G, Namani T, Hofmann H. Slow domain reconfiguration causes power-law kinetics in a two-state enzyme. Proc Natl Acad Sci USA 115, 513–518 (2018).

44. Lasitza-Male T, et al. Membrane Chemistry Tunes the Structure of a Peptide Transporter. Angew Chem Int Ed Engl 28, 51 (2020).

45. Soranno A, et al. Quantifying internal friction in unfolded and intrinsically disordered proteins with single molecule spectroscopy. Proc Natl Acad Sci USA 109, 17800–17806 (2012).

46. Soranno A, Zosel F, Hofmann H. Internal friction in an intrinsically disordered protein-Comparing Rouse-like models with experiments. J Chem Phys 148, 123326 (2018).

47. Rouse PE. A Theory of the Linear Viscoelastic Properties of Dilute Solutions of Coiling Polymers. J Chem Phys 21, 1272–1280 (1953).

48. Stappert J, Kemler R. A short core region of E-cadherin is essential for catenin binding and is highly phosphorylated. Cell Adhes Commun 2, 319–327 (1994).

49. Kramers H. Brownian motion in a field of force and the diffusion model of chemical reactions. Physica 7, 284–304 (1940).

50. Zwanzig R. Diffusion in a rough potential. Proc Natl Acad Sci USA 85, 2029–2030 (1988).

51. Frauenfelder H, Sligar SG, Wolynes PG. The energy landscapes and motions of proteins. Science 254, 1598–1603 (1991).

52. Ansari A, et al. Protein states and proteinquakes. Proc Natl Acad Sci USA 82, 5000–5004 (1985).

53. Wensley B, et al. Experimental evidence for a frustrated energy landscape in a three-helix-bundle protein family. Nature 463, 685–U122 (2010).

54. Fuxreiter M, Tompa P. Fuzzy Complexes: A More Stochastic View of Protein Function. Springer US (2012).

55. Bigman LS, Levy Y. Tubulin tails and their modifications regulate protein diffusion on microtubules. Proc Natl Acad Sci USA 117, 8876–8883 (2020).

56. Sheu S-Y, Yang D-Y, Selzle HL, Schlag EW. Energetics of hydrogen bonds in peptides. Proc Natl Acad Sci USA 100, 12683–12687 (2003).

57. Sievers S, Fritzsch C, Grzegorczyk M, Kuhnen C, Müller O. Absolute beta-catenin concentrations in Wnt pathway-stimulated and non-stimulated cells. Biomarkers 11, 270–278 (2006).

58. Theillet F-X, et al. Physicochemical properties of cells and their effects on intrinsically disordered proteins (IDPs). Chem Rev 114, 6661–6714 (2014).

59. Choi H-J, Loveless T, Lynch AM, Bang I, Hardin J, Weis WI. A conserved phosphorylation switch controls the interaction between cadherin and β-catenin in vitro and in vivo. Dev Cell 33, 82–93 (2015).

60. Erijman A, Dantes A, Bernheim R, Shifman JM, Peleg Y. Transfer-PCR (TPCR): a highway for DNA cloning and protein engineering. J Struct Biol 175, 171–177 (2011).

61. Unger T, Jacobovitch Y, Dantes A, Bernheim R, Peleg Y. Applications of the Restriction Free (RF) cloning procedure for molecular manipulations and protein expression. J Struct Biol 172, 34–44 (2010).

62. Frey S, Goerlich D. Purification of protein complexes of defined subunit stoichiometry using a set of orthogonal, tag-cleaving proteases. J Chromatogr A 1337, 106–115 (2014).

63. Müller BK, Zaychikov E, Bräuchle C, Lamb DC. Pulsed interleaved excitation. Biophys J 89, 3508–3522 (2005).

64. Kapanidis AN, Laurence TA, Lee NK, Margeat E, Kong X, Weiss S. Alternating-laser excitation of single molecules. Acc Chem Res 38, 523–533 (2005).

65. Eggeling C, et al. Data registration and selective single-molecule analysis using multi-parameter fluorescence detection. J Biotechnol 86, 163–180 (2001).

66. Hoffmann A, et al. Mapping protein collapse with single-molecule fluorescence and kinetic synchrotron radiation circular dichroism spectroscopy. Proc Natl Acad Sci USA 104, 105–110 (2007).

67. Schuler B. Application of single molecule Förster resonance energy transfer to protein folding. Methods Mol Biol 350, 115–138 (2007).

68. Hillger F, Nettels D, Dorsch S, Schuler B. Detection and analysis of protein aggregation with confocal single molecule fluorescence spectroscopy. J Fluoresc 17, 759–765 (2007).

69. Benke S, Nettels D, Hofmann H, Schuler B. Quantifying kinetics from time series of single-molecule Förster resonance energy transfer efficiency histograms. Nanotechnology 28, 114002 (2017).

70. Haynes WM. Handbook of Chemistry and Physics, 95 edn. CRC Press (2014).

71. Pace CN. Determination and analysis of urea and guanidine hydrochloride denaturation curves. METHOD ENZYMOL 131, 266–280 (1986).

72. Schuler B, Lipman E, Eaton W. Probing the free-energy surface for protein folding with single-molecule fluorescence spectroscopy. Nature 419, 743–747 (2002).

73. Dertinger T, Pacheco V, von der Hocht I, Hartmann R, Gregor I, Enderlein J. Two-focus fluorescence correlation spectroscopy: a new tool for accurate and absolute diffusion measurements. Chemphyschem 8, 433–443 (2007).

74. Wilkins D, Grimshaw S, Receveur V, Dobson C, Jones J, Smith L. Hydrodynamic radii of native and denatured proteins measured by pulse field gradient NMR techniques. Biochemistry 38, 16424–16431 (1999).

75. Aznauryan M, Nettels D, Holla A, Hofmann H, Schuler B. Single-molecule spectroscopy of cold denaturation and the temperature-induced collapse of unfolded proteins. J Am Chem Soc 135, 14040–14043 (2013).

76. Nettels D, et al. Single-molecule spectroscopy of the temperature-induced collapse of unfolded proteins. Proc Natl Acad Sci USA 106, 20740–20745 (2009).

77. Russinova E, Tretyachenko-Ladokhina V, Vele OE, Senear DF, Alexander Ross JB. Alexa and Oregon Green dyes as fluorescence anisotropy probes for measuring protein– protein and protein–nucleic acid interactions. Anal Biochem 308, 18–25 (2002).

78. Likhachev ER. Dependence of water viscosity on temperature and pressure. Tech Phys 48, 514–515 (2003).

79. O’Brien EP, Morrison G, Brooks BR, Thirumalai D. How accurate are polymer models in the analysis of Förster resonance energy transfer experiments on proteins? J Chem Phys 130, 124903 (2009).

80. Sanchez I. Phase Transition Behavior of the Isolated Polymer Chain. Macromolecules 12, 980–988 (1979).

81. Ziv G, Haran G. Protein folding, protein collapse, and tanford’s transfer model: lessons from single-molecule FRET. J Am Chem Soc 131, 2942–2947 (2009).

82. Sherman E, Haran G. Coil-globule transition in the denatured state of a small protein. Proc Natl Acad Sci U S A 103, 11539–11543 (2006).

83. Zheng W, Zerze GH, Borgia A, Mittal J, Schuler B, Best RB. Inferring properties of disordered chains from FRET transfer efficiencies. J Chem Phys 148, 123329 (2018).

84. Gopich IV, Nettels D, Schuler B, Szabo A. Protein dynamics from single-molecule fluorescence intensity correlation functions. J Chem Phys 131, 095102 (2009).

85. Dignon GL, Zheng W, Kim YC, Best RB, Mittal J. Sequence determinants of protein phase behavior from a coarse-grained model. PLoS Comput Biol 14, e1005941 (2018).

86. Debye P, Hückel E. Zur Theorie der Elektrolyte: I. Gefrierpunktserniedrigung und verwandte Erscheinungen. Phys Z 24, 185–206 (1923).

87. Ashbaugh HS, Hatch HW. Natively unfolded protein stability as a coil-to-globule transition in charge/hydropathy space. J Am Chem Soc 130, 9536–9542 (2008).

88. Kapcha LH, Rossky PJ. A simple atomic-level hydrophobicity scale reveals protein interfacial structure. J Mol Biol 426, 484–498 (2014).

89. Plimpton S. Fast Parallel Algorithms for Short-Range Molecular Dynamics. J Comput Phys 117, 1–19 (1995).

90. Anderson JA, Glaser J, Glotzer SC. HOOMD-blue: A Python package for high-performance molecular dynamics and hard particle Monte Carlo simulations. Comput Mater Sci 173, 109363 (2020).

91. McCarney ER, et al. Site-specific dimensions across a highly denatured protein; a single molecule study. J Mol Biol 352, 672–682 (2005).

92. Makarov DE. Spatiotemporal correlations in denatured proteins: The dependence of fluorescence resonance energy transfer (FRET)-derived protein reconfiguration times on the location of the FRET probes. The Journal of chemical physics 132, 035104 (2010).

